# High Coverage Mitogenomes and Y-Chromosomal Typing Reveal Ancient Lineages in the Modern-day Székely Population in Romania

**DOI:** 10.1101/2022.11.07.515481

**Authors:** Noémi Borbély, Orsolya Székely, Bea Szeifert, Dániel Gerber, István Máthé, Elek Benkő, Balázs Gusztáv Mende, Balázs Egyed, Horolma Pamjav, Anna Szécsényi-Nagy

## Abstract

Here we present 115 whole mitogenomes and 92 Y-chromosomal STR and SNP profiles from a Hungarian ethnic group, the Székelys (in Romanian: Secuii, in German: Sekler) living in southeast Transylvania (Romania). The Székelys can be traced back to the 12th century in the region, and numerous scientific theories exist as to their origin. We carefully selected sample providers that had local ancestors inhabiting small villages in the area of Odorheiu Secuiesc/Székelyudvarhely in Romania. The results of our research and the reported data signify a qualitative leap compared to previous studies, since complete mitochondrial DNA sequences and Y-chromosomal data containing 23 STRs have not been available from the region until now. We evaluated the results with population genetic and phylogenetic methods, in the context of the modern and ancient populations that are either geographically or historically related to the Székelys. Our results demonstrate a predominantly local uniparental make-up of the population that also indicates limited admixture with neighbouring populations. Phylogenetic analyses confirmed the presumed eastern origin of certain maternal (A, C, D) and paternal (Q, R1a) lineages and, in some cases, they could also be linked to ancient DNA data from Migration Period (5^th^-9^th^ centuries AD) and Hungarian Conquest Period (10th century AD) populations.

## Introduction

The Székelys (also known as Szeklers or Seklers) are a Hungarian-speaking minority that has been living in Transylvania (Romania) for more than 800 years. Several theories have been elaborated about the origin of the Székelys over time, but it is still an unresolved question to this day. They have been identified as descendants of Hunnic, Avar, Kabar, Volga Bulgarian (Onogur) and Hungarian ethnic groups. The story of their Hunnic origin was elaborated by mediaeval Hungarian chroniclers (who, by doing so, increased the authority of the Árpád dynasty and created the legal basis for the Hungarian conquest), so the Székelys’ own Hunnic ‘tradition’ seems to have developed secondarily as a result of this and, due to the lack of evidence, modern historiography and archaeology have not considered the Székelys to be of Hunnic origin for some time [1]. According to some scholars, the ancestors of the Székelys were ethnic groups separated from the Volga Bulgarians, and were thus of Turkish origin. According to this idea, the accession of these Bulgarian tribes to the Hungarians would have taken place even before the Hungarian conquest of the Carpathian Basin in 895 AD. The theory, however, that attempted to connect the Székely folk name with the Askal/Äskäl tribe of the Bulgarians turned out to be linguistically incorrect [2]. There are scholars who consider the Székelys to be the remnants of the late Avar population who, according to the scholars’ assumption, spoke the Hungarian language. Other scholars assume that the Székelys were originally Hungarian ethnic groups who guarded the various border sections of the early Hungarian Kingdom, primarily at the western ends, and later, in the 12th–13th centuries, the majority of them were resettled in Transylvania in order to stop the Cuman and later Tatar incursions that threatened the eastern borders [3]. The first written mention of the Székelys originates from the 12th century, mentioning them as military auxiliaries of the Hungarians along the Pechenegs, then still in the western border region.

At the moment, the research faces a serious contradiction. On the one hand, we see a Székely population with its own name and traditions, which, after observing typical military service to the king, may seem like an ‘auxiliary people’ who joined the Hungarians, whose territorial organisation was not the usual county of the rest of the Hungarians, but the district/*sedes* typical of foreign ethnic groups; the seemingly distant connections of this population towards Asia and the late Avar period have recently been raised by physical anthropological research [4]. On the other hand, their archaeological findings from the Árpád period do not differ from the findings of the rest of the Hungarian population and do not show special oriental features, just as their Hungarian dialect does not indicate a change of language and the subsequent acquisition of the Hungarian language. Their placement in small patches at critical points of the country’s border reflects the conscious organisation of the early Árpádian kings, for the sake of border and land protection. Their significant mediaeval privileges were based on their continuous military service, which made them important actors of the time, and this continued in the early modern period, in the territory of the independent Principality of Transylvania. To this day, their mother tongue is the clearly spoken Hungarian.

A branch of the Hungarian ethnic group known as the Csángó is also related to the Székelys, the Csángós of Ghimeş/Gyímes, who moved from the area of Ciuc/Csík district to the valley of the Trotuş/Tatros river on the border of Transylvania and Moldavia from the early modern period (or perhaps earlier) and whose language is thus closely related to the Csík dialect [5]. Due to their close relationship with the Székelys, the Gyímes csángós are also analysed in this study from published sources. They should not be confused with other Csángó groups living in other areas of Moldavia and speaking a different dialect, whose origins are also different.

In recent decades, molecular genetics studies have described the genetic make-up of some of the urban Székely (Miercurea Ciuc/Csíkszereda and Corund/Korond) and Ghimeş Csángó groups, investigating maternally inherited mitochondrial DNA (mtDNA) [6–8] and paternal Y-chromosomal [9–11] lineages. Most of these studies lacked thoroughly planned and executed sample collection; thus, we cannot be sure that all sample donors had local ancestors. The studies revealed an increased number of Central or Eastern Asian lineages in the Székely population compared to other Hungarian-speaking populations. In addition to former uniparental studies, genome-wide genotype data from 24 Székely individuals from the commune of Corund were analysed and compared with the genotype data of Hungarians [12]. The genetic research on the Székely population does not currently have databases containing complete mitochondrial genomes, that would have been based on accurate sampling. Both of these have great importance in evaluating genetic continuity between present-day and ancient populations.

Besides Csángós and Székelys, the population of Hungary has also been investigated [13]. Hungarian paternal lineages from Hungary were reported by Völgyi et al. in 2009 and by Pamjav et al. in 2017 [14,15]. In the latter study, all samples were collected in the Bodrogköz, which is a geographical area in the Upper-Tisza region in north-eastern Hungary bordered by the Bodrog and Tisza rivers. Due to its isolated nature, the dataset from Bodrogköz has been treated separately from the Hungarian data in our study. The authors assumed that its present-day population is likely to preserve ancient markers and lineages, as its former inhabitants had a better chance of surviving both Tatar (Mongol) and Ottoman invasions than groups living in some of the other affected regions [15]. There is scarce genetic data available on the Romanian population—only one mitochondrial DNA sampling has been reported in Romania, and haplogroup results, which are based on mtDNA control region and coding marker data, were made accessible by Cocos et al. in 2017 [16].

In this study, we aimed to reconstruct the uniparental gene pool of the Székely population that existed 100–150 years ago by finding elderly sample donors living in isolated villages and carefully documenting their genealogies. Furthermore, we aimed to monitor any regional genetic structure discrepancies of the Hungarian-speaking population and to confirm preliminary uniparental genetic studies that revealed an increased number of Eastern Eurasian lineages in isolated populations, compared to populations of larger cities nearby. We present new genetic data containing 115 newly sequenced whole mitochondrial genomes, and 92 Y-chromosomal Short Tandem Repeat (STR) haplotypes and haplogroups of a Székely population that has not been sampled before, and compare them to recent Eurasian and available ancient DNA (aDNA) data to gain further knowledge about their genetic history.

## Material and Methods

### DNA samples, extraction, amplification and sequencing

Samples were collected with buccal swabs by researchers from ELRN RCH Institute of Archaeogenomics, the ELTE University of Budapest and the Sapientia Hungarian University of Transylvania. The samples were taken from 115 mostly unrelated individuals of the Székely population of Transylvania, Romania. The selected individuals spoke Hungarian as their mother tongue and had Hungarian surnames. All sampled individuals agreed and gave their written consent to the anonymous use of their samples in this study. Their ancestors had been documented for two generations, and these ancestors were born in the same region of Transylvania and had declared themselves as Székelys. The following villages in Udvarhely county were included in the sampling, near the town Odorheiu Secuiesc, which has appeared in written records from the 14^th^ century onward [17]: Inlăceni/Énlaka, n = 9; Firtănuş/Firtosmartonos, n = 7; Ulieş/Kányád, n = 21; Mugeni/Bögöz, n = 13; Goagiu/Gagy, n = 11; Avrămeşti/Szentábrahám, n = 13; Cecheşti/Csekefalva, n = 9; Dobeni/Székelydobó, n = 12; Văleni/Patakfalva, n = 7; Forțeni/Farcád n = 13 (Figure 1, Supplementary Table S1).

**Figure 1.**
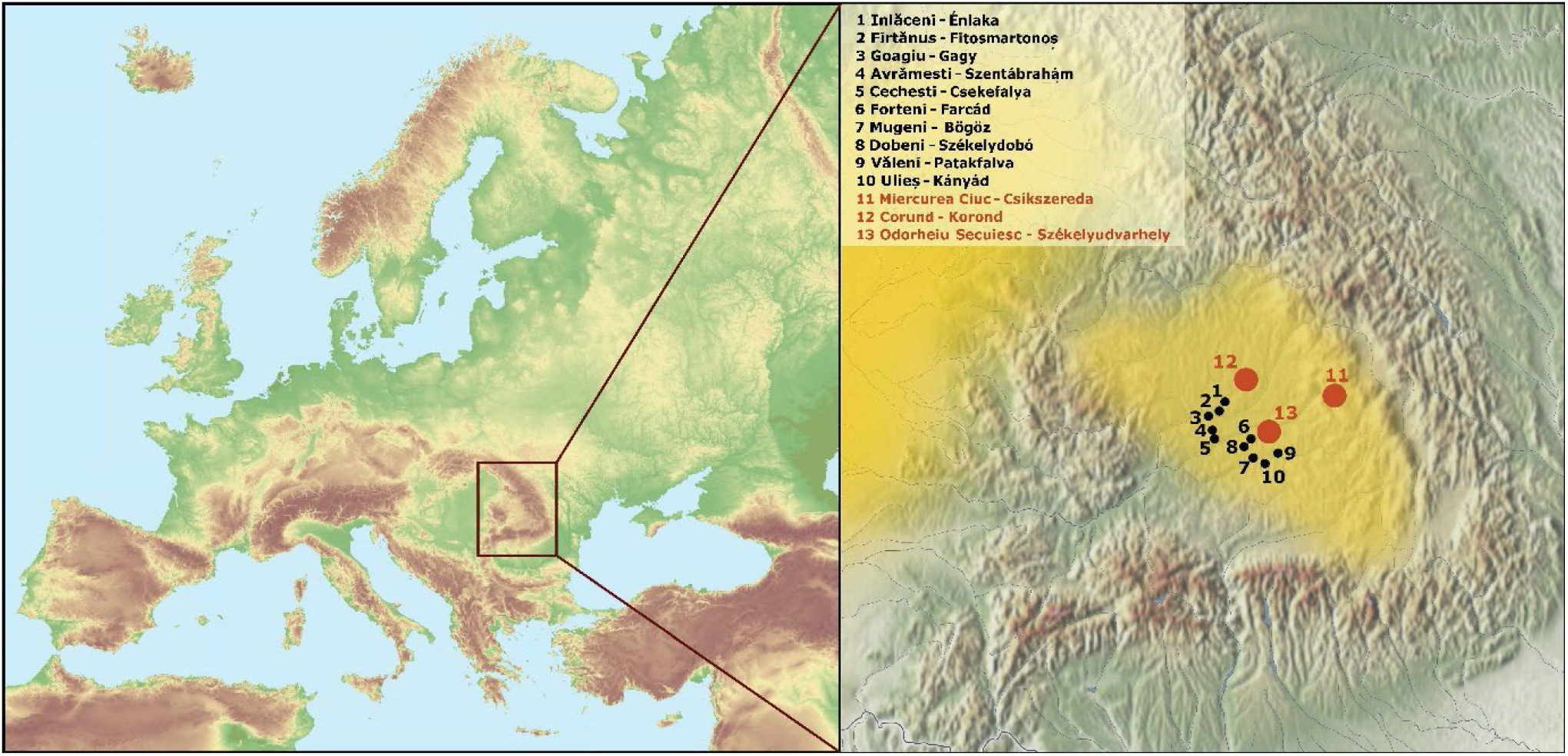
Map of Europe and the Transylvanian part of Romania showing the Székely villages where the DNA samples were collected (in black). The yellow shadings indicate the settlement areas of the Hungarian-speaking populations, including the Székelys. Red circles indicate previously collected and published Székely datasets [6], [8] and the city of Odorheiu Secuiesc. The map of Europe was downloaded from MAPSWIRE [18], licensed under CC BY 4.0. The map of the Carpathian Basin is owned by the Institute of Archaeogenomics, Research Centre for the Humanities, Eötvös Loránd Research Network; modifications were made in Adobe Acrobat Pro DC and Inkscape 1.1.1.

DNA was extracted with QIAamp DNA Mini Kit (Qiagen) according to the producer’s buccal swab spin protocol. The concentration of the samples was measured with Qubit™ dsDNA High Sensitivity Assay Kit (Thermo Fisher Scientific).

The amplification of the whole mtDNA was performed with the Expand™ Long Range dNTPack kit (Sigma Aldrich) according to Fendt et al. [19] (primer sequence 5’–3’, forward ‘A’ (FA): AAATCTTACCCCGCCTGTTT; reverse ‘A’ (RA): AATTAGGCTGTGGGTGGTTG; forward ‘B’ (FB): GCCATACTAGTCTTTGCCGC; reverse ‘B’ (RB): GGCAGGTCAATTTCACTGGT). We amplified the mtDNA in two fragments, and modified the PCR program according to the length of the fragments. Conditions used for long-range PCR consisted of an initial denaturation step of 2 min at 92°C followed by 10 cycles of denaturation at 92°C for 10 s, annealing at 60°C for 15 s and elongation at 68°C for 8 m 30 s, 10 cycles of denaturation at 92°C for 10 s, annealing at 60°C for 15 s and elongation at 68°C for 8 m 50 s, 15 cycles of denaturation at 92°C for 10 s, annealing at 60°C for 15 s and elongation at 68°C for 9 m 10 s, and a final elongation step at 68°C for 7 min. The amplification reaction was checked on 0.8% agarose gel and visualised after EcoSafe staining with UV transillumination. We pooled the two separately amplified fragments, then purified the amplicons with the QIAquick PCR Purification Kit (Qiagen). The concentration of the PCR products was measured with Qubit™ dsDNA Broad Range Assay Kit (Thermo Fisher Scientific).

NEBNext Ultra II FS DNA Library Prep Kit for Illumina kit was used for the preparation of the mtDNA libraries. The products were checked with the Agilent D1000 ScreenTape Assay on the 4200 Tapestation system. Next-generation sequencing was performed on the Illumina Miseq System (Illumina) using Illumina Miseq Reagent Kit V2 (2 × 150 cycles) sequencing kit. Indexed libraries’ final concentrations were adjusted to 4 nM. Samples were pooled together with regard to the calculated desired coverage. Five percent of PhiX was used to increase the heterogeneity of samples.

We analysed 92 male samples in the Laboratory of Reference Sample Analysis of the Department of Genetics, Directorate of Forensic Expertise, Hungarian Institute for Forensic Sciences in Budapest. DNA was surveyed for STR variation using the Promega PowerPlex Y23 for the Székely population, including 23 Y-STR loci. ABI3130 Genetic Analyzer and GeneMapper ID-X v.1.2 software were used for fragment analyses of PCR products. The results of the Y-chromosomal Short Tandem Repeat (STR) analyses were verified by haplogroup-defining Single Nucleotide Polymorphism (SNP) markers (see Supplementary Table S2) on ABI 7500 Real-time PCR instrument using SDS.1.2.3 software.

### Pre-processing of the sequencing data

A custom in-house analytic pipeline was applied on the Illumina sequencing data [20]. Pairedend reads were merged together with SeqPrep master [21]. At a maximum of one mismatch, the one base with higher base quality was accepted and the overlapping reads with two or more mismatches were discarded. The pre-processed reads were mapped to the rCRS reference sequence using BWA v.0.7.5 [22] with MAPQ of 30. Samtools v.1.3.1 [23] was used for further data processing, such as indexing, removing PCR duplications and creating bcf files.

Mitochondrial haplogroup determinations were performed by HaploGrep2 [24], which uses Phylotree mtDNA tree Build 17 [25,26].

Y-haplogroups were assigned based on Y-STR data using nevgen.org, as well as based on Y-SNP genotyping by TaqMan assay on a Real-time PCR platform. Terminal Y-SNPs were verified on the Y tree of ISOGG version 15.34 [27].

We created and visualised the median-joining (MJ) network of the whole mitochondrial genomes of our dataset with the PopArt program [28]. The input file of PopArt was made by DnaSP [29].

### Phylogenetic analysis of the mtDNA

We used all publicly available data to compile datasets for each mtDNA haplogroup with sequences that belonged to the same haplotype or were closely related to it. For each phylogenetic tree, 50–150 mtDNA sequences were used. For every haplotype, all available closely related mitochondrial genome sequences in NCBI were downloaded. Multiple alignments for each haplogroup were performed with ClustalO within SeaView [30]. The alignments were checked and corrected manually where necessary. Next, neighbour-joining (NJ) trees were generated by PHYLIP version 3.6 [31]. The phylogenetic trees were then drawn using Figtree version 1.4.2 [32].

Most of the data used for the neighbour-joining mitochondrial phylogenetic trees are from the NCBI database; IDs and sources of other data are available in Supplementary Tables S5.

### Population genetic analysis

Principal component analysis was performed based on the mtDNA haplogroup frequencies of 56 modern and two ancient populations. In the PCA of the modern populations, we considered 36 mitochondrial haplogroups. The PCAs were carried out using the prcomp function in R v4.0.0. [33] and visualised in two-dimensional plots with two principal components (PC1 and PC2 or PC1 and PC3).

For a Ward-type hierarchical clustering we involved the same population datasets as for PCAs. Based on the mtDNA haplogroup frequencies, we calculated PC-scores in R v4.0.0 [33], then applied PC1–PC6 scores using the Euclidean distance measurement and ward.D method. We visualised the results as a dendrogram with the hclust library.

To calculate the inter-population variability of the mtDNA genetic profiles characteristic of the three Székely populations, we performed analysis of molecular variance (AMOVA) using Arlequin v3.5.2.2 software [34].

We calculated population pairwise FST and linearised Slatkin F_ST_ values based on the whole mitochondrial genome sequences of 3981 modern-day individuals (classified into 21 groups) and 362 ancient individuals (classified into 7 groups) using Arlequin v3.5.2.2. [34] with the following settings: Tamura & Nei substitution model with 10,000 permutations, significance level of 0.05, gamma value of 0.3.

We used the same linearised Slatkin F_ST_ values for clustering in Python using the seaborn [35] clustermap function (parameters: metric = ‘correlation’, method = ‘complete’).

### Analysis of Y-STR variations

MJ networks were constructed using Network v10.1.0.0 and the results were visualised with Network Publisher v2.1.2.5 [36]. The following settings were used in Network v10.1.0.0: Network Calculation: Median-Joining [37], Optional Post-processing: MP calculation [38] (selected option: Network containing all shortest trees, and list of some of the shortest trees sufficient to generate the network) and in Network Publisher, shortest tree visualisation was applied and coloured according to the haplogroups and sample provenance.

## Results

The sample pool of this study consisted of 92 male and 23 female participants, all Székely individuals from Transylvania, Romania (for detailed information see Supplementary Table S1). We performed whole mitochondrial genome enrichment and next-generation sequencing (NGS) to obtain 115 complete mitogenomes. In addition, we investigated the Y-STR profiles (23 STRs) and Y-SNP data of the 92 male individuals (see Supplementary Table S2).

### Maternal lineages in the dataset

#### Haplogroup-based analyses

115 high-coverage mitochondrial genomes were obtained with NGS methods (from 111.46x to 276.83x coverage), with a mean coverage of 233.56x. The 115 complete mitochondrial genomes were classified into 72 different haplotypes. These mitochondrial haplotypes are mainly present in European regions, but there were several haplotypes predominantly found in present-day Asian or Near Eastern populations. The new dataset consisted of the following macro haplogroups: A, C, D, H, HV, HV0, I, J, K, T, U, N, R, V, W and X. A list of the mtDNA sub-haplogroups found in the Székely population is in Table 1. The overall haplogroup composition of the investigated Székely population was similar to the formerly described Székely populations in Miercurea Ciuc and Corund, and to the Hungarian population in Hungary as well [7,8]. Around Odorheiu Secuiesc, most of the individuals belonged to haplogroup H—as expected in a European population [39], and as had been observed in the case of earlier studied Székely, Hungarian and Csángó populations (see Figure 2). Compared to the populations of Miercurea Ciuc and Corund, some differences were conspicuous. We observed a higher proportion of haplogroups I, T2, HV and W, and a lower proportion of haplogroup K than previous studies; furthermore, no T1 was present in our dataset. The proportion of the Eastern Eurasian-derived haplogroups was higher in the population of Miercurea Ciuc than around Odorheiu Secuiesc and in Corund. The Csángó population from Ghimeş stood out slightly in the comparison due to its high proportion of the haplogroup K and lower proportion of H [7]. All three Székely populations had a significant proportion of mitochondrial haplogroups with Eastern Eurasian prevalence (groups A, B, C, D, G and Y).

**Figure 2.**
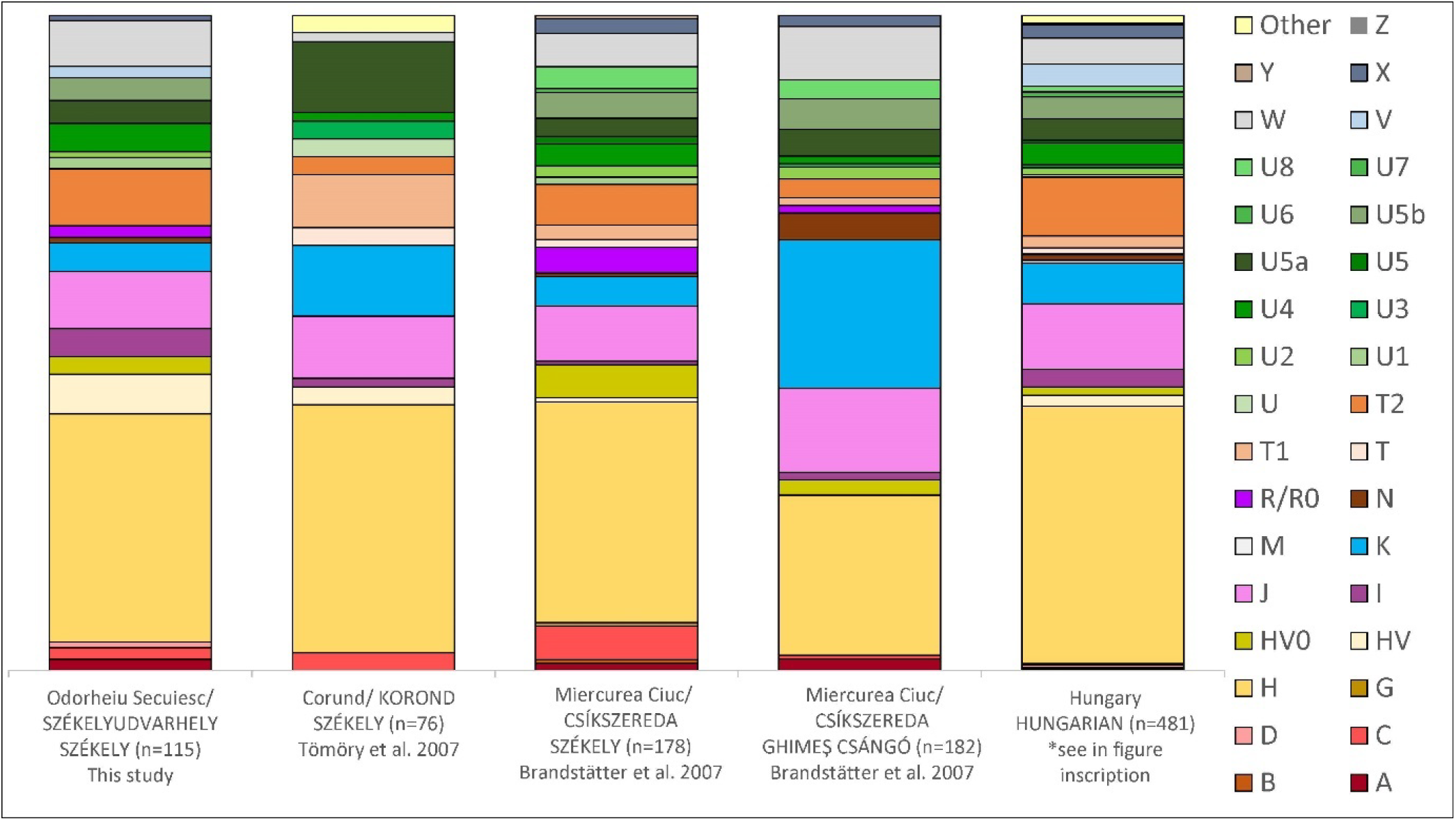
Haplogroup composition of the investigated Székely population around Odorheiu Secuiesc (n = 115) compared to other Székely populations [7,8], the Csángó population [7] and Hungarians living in Hungary (n = 481).

**Figure 3.**
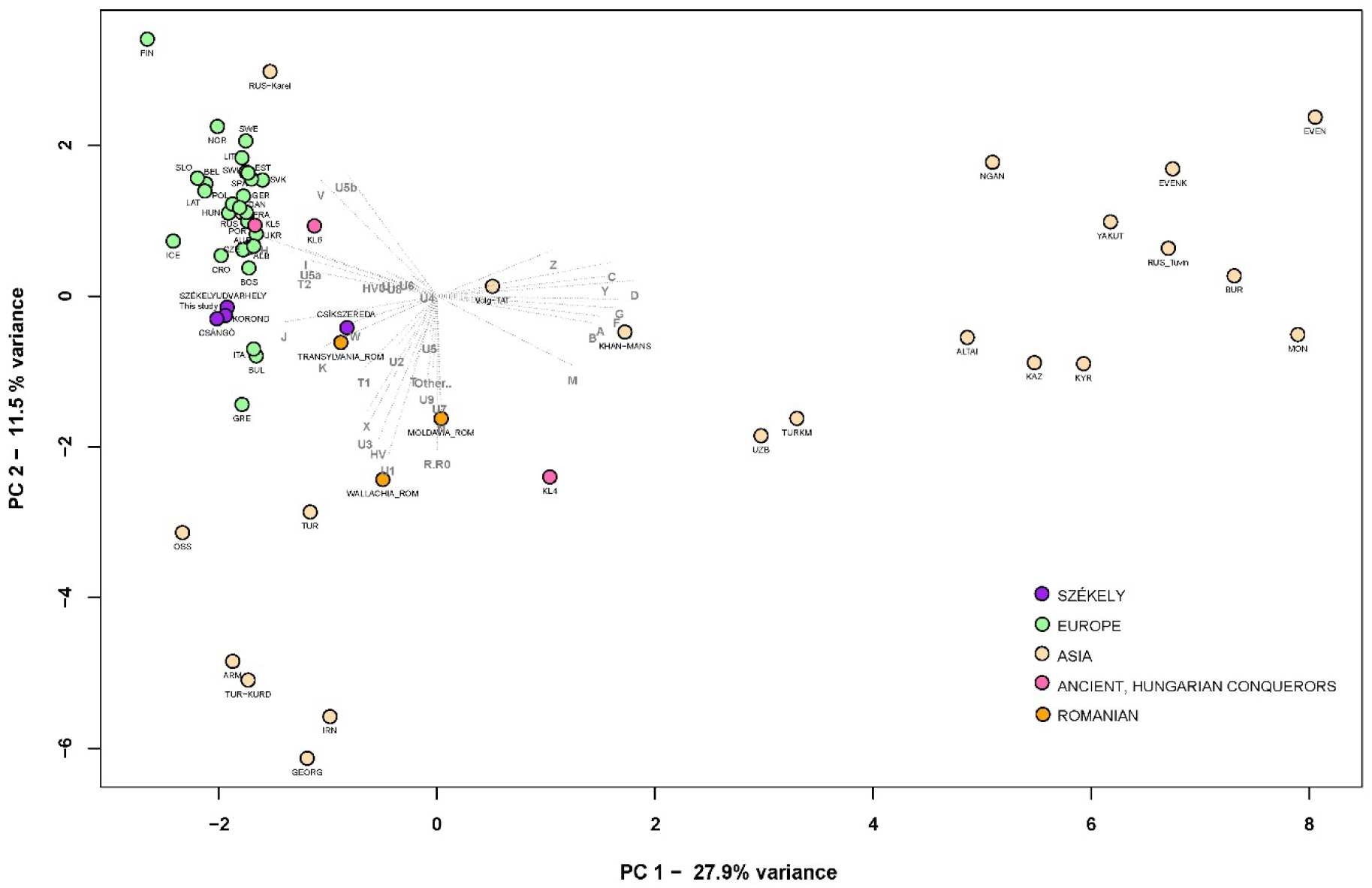
PCA plot with 56 modern and three ancient populations (36,803 samples), representing first and second principal components (39.4% of the total variance): PCA analysis based on haplogroup frequencies in Eurasian modern populations and three ancient populations. The selected ancient populations are the Hungarian Conquest Period (10th century AD) populations of the Carpathian Basin [41–44]. KL4-5-6 groups indicate different cemetery types in the Hungarian Conquest Period, as used in Szeifert et al. 2022 [45]. The investigated Székely population and previously examined Székely groups are marked in purple, the ancient populations from Hungary in pink, modern-day Romanian populations in orange, other modern-day Europeans in green and Asian populations in beige. The PCA shows a clear separation of Eastern (right side of the plot) and Western (left side of the plot) populations. For further information see Supplementary Table S3.

**Table 1.** Modern-day distribution peaks of the mtDNA sub-haplogroups found in the Székely population based on the IanLogan database [40]. Column ‘n’ gives the absolute frequency of the given haplogroup in the studied Székely population around Odorheiu Secuiesc/Székelyudvarhely.

| Observed mtDNA haplogroup in this study | n | Frequency in the Székely population | Modern-day presence/geographic distribution peaks of the haplogroups |
| --- | --- | --- | --- |
| **A+152+16362** | 1 | 0.87% | Eastern Asia (Han Chinese, Japanese) |
| **A12a** | 1 | 0.87% | East-Asia (Evenk, Yakut, Mansi) |
| **C4a1a3** | 1 | 0.87% | Northeastern-Asia, Central Siberia (Even, Evenk, Yukaghir, Yakut, Altai-Kizhi, Buryat) |
| **C5c1a** | 1 | 0.87% | Middle East (Iranian), Europe |
| **D4e4** | 1 | 0.87% | Northeastern-Asia, Central Siberia (Tungusic, Evenk, Pamir, Uyghur, Buryat) |
| **H and subclades (H, H1, H1a3b, H1b1a, H1c, H1c1b, H1c20, H1q, H1u2, H24a, H26a1, H2a1a, H2a3, H3, H3h, H4a1, H5, H5'36, H55, H5a1, H6a1a, H11a1, H13a1c)** | 39 | 33.91% | Europe |
| **HV, HV0 and subclades (HV+16311, HV6, HV11, HV0a1, HV0f)** | 10 | 8.70% | Caucasus (Armanians), Middle East (Turks, Iranians), Europe |
| **I2** | 1 | 0.87% | Northern Europe (English, Irish, Norwegian), Caucasus |
| **I5 and subclades (I5a1, I5a4)** | 4 | 3.48% | Europe, Middle East (Turks) |
| **J and subclades (J1b1a1, J1c2, J1c2p, J1c10, J2b1)** | 10 | 8.70% | Europe (Finnish, English), Middle East (Iranian) |
| **K and subclades (K1a+195, K1a2, K1b2a2, K1c1a, K2b1a4)** | 5 | 4.35% | Europe, Eastern Europe (Russian), Baikal Region, Caucasus |
| **N1b1a** | 1 | 0.87% | Caucasus, Middle East, Eastern Europe |
| **R0a2** | 3 | 2.61% | Middle East (Iranians), Arabian Peninsula, Horn of Africa |
| **T, T2 and subclades (T2a1b1ba1, T2b, T2b25, T2b4, T2f1a1, T2g1a)** | 10 | 8.70% | Europe, Caucasus, Central Siberia, Middle East |
| **U subclades (U1a1a1, U1a1a1a, U2e1a1, U4a, U4a2, U5a1a1, U5a1b1a, U5b1a, U5b2b1b)** | 16 | 13.91% | Middle East (Iranian), Europe, Russia |
| **V and subclade (V, V1a1b)** | 2 | 1.74% | Europe (Finnish), Eastern Europe (Volga Tatars) |
| **W and subclades (W, W1e1a, W1i, W3a1d, W5a1a)** | 8 | 6.96% | Europe, Europe, North-Caucasus, Central-Asia, India, Iran |
| **X2e1** | 1 | 0.87% | Europe |

We used Principal Component Analysis (PCA) in order to comparatively visualise the mtDNA haplogroup composition of the different populations (see Supplementary Table S3). The investigated Székely population (Odorheiu Secuiesc) was positioned among European populations, closest to other Székelys, Csángós, Croatians, Bosnians, modern Czech populations and Transylvanian Romanians. It was not possible to further examine all connections at the complete mitogenome sequence level due to the lack of whole mitogenome data in some populations.

Data on the PCA were also displayed using the hierarchical ward-clustering method (see Supplementary Information Figure S1). The clustering confirms the connection of the studied Székely group with Europeans, but also separates the Miercurea Ciuc, Ghimeş Csángó and Corund groups from Odorheiu Secuiesc. This difference from the PC1-2 plot is also visible on the PC3, where the Odorheiu Secuiesc group becomes distant from the others (Supplementary Information Figure S2).

Since the other two Székely groups were only analysed for Hypervariable Region I of the mitochondrial genome, the sequence-based comparison of the three groups with limited conclusions is discussed in the Supplementary Information.

#### Whole mitogenome sequence-based evaluations

##### F_ST_ analyses

We analysed the whole mitogenomes (16,569 base pairs) at the DNA sequence level, and calculated Slatkin F_ST_ values (see Supplementary Table S4). A heatmap with clustering of F_ST_ values was created to visualise the genetic differentiation of the examined populations (Supplementary Information Figure S3). The Székelys cluster on the European branch with Hungarians, but the Serbians and the Conquest Period Hungarians are the most similar to them.

Whole mitochondrial data are missing from Romania, and the Slovakian and Czech datasets are also limited; therefore, the resolution of that analysis is restriced. Among the ancient populations, the KL6 group, which was discussed by Szeifert et al. as comprising large village cemeteries opened in the 10th century and used until the 11th and 12th centuries in the Hungarian Kingdom [45], was the closest to the Székelys.

##### Phylogenetic analyses of the Székely maternal lineages

The median-joining network of the Székely mitogenomes showed the variable distribution of the maternal lineages among the sampled villages (see Figure 4). Most of the haplogroups were shared among the villages, and shared lineages (haplotypes) were also found among certain H, T, U4, U5 and W haplotypes.

**Figure 4.**
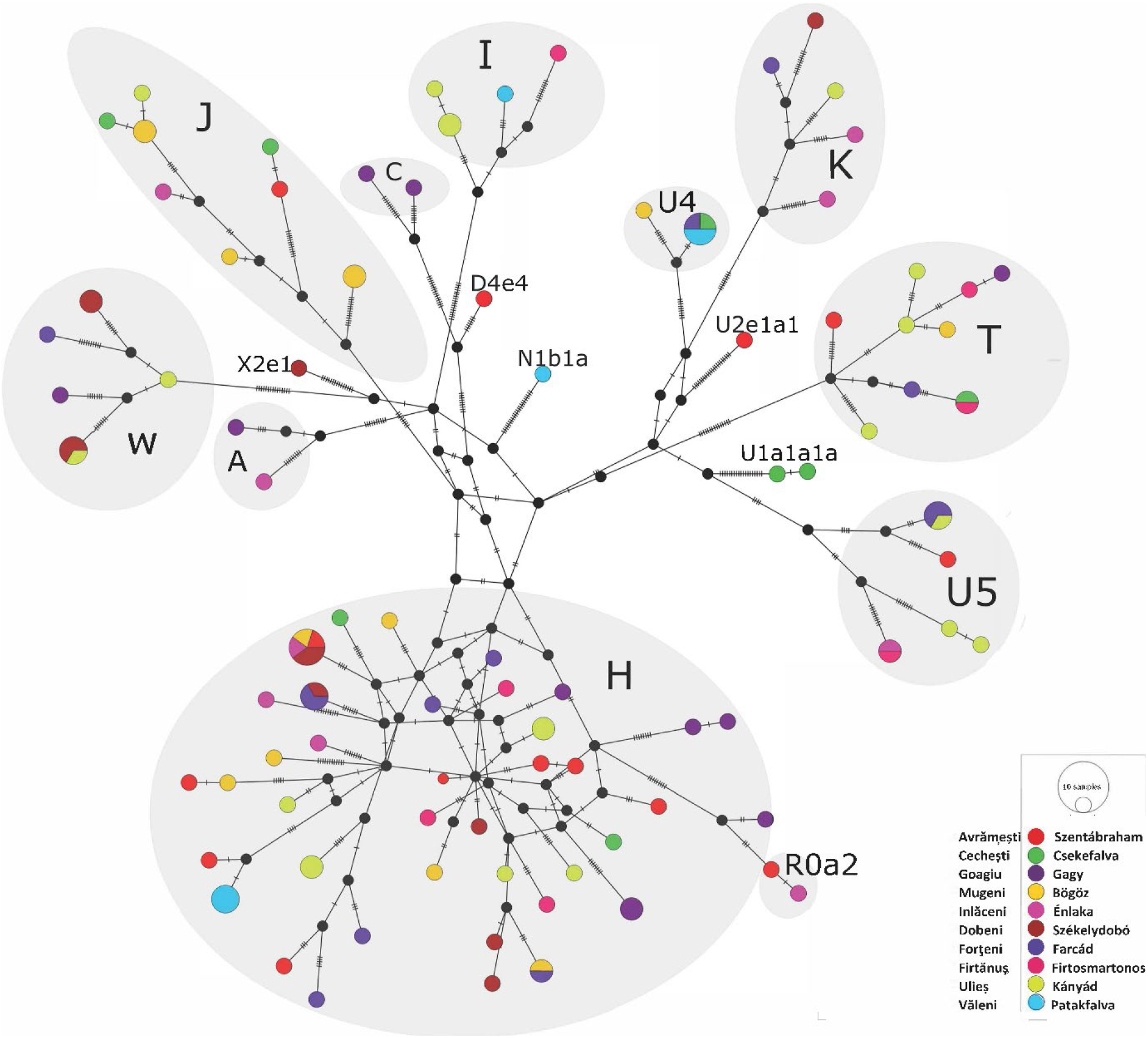
Median-joining network of 115 modern-day Székely mitogenomes. The sequences contained 484 variable sites and belonged to 72 haplotypes. The figure was created with the PopArt program.

Three individuals belonging to the described Eastern Eurasian haplogroups originated from the same village (Goagiu), although we detected Eastern Eurasian haplogroup types in the Avrǎmeşti and Inlǎceni villages as well.

Since analyses of mitogenomes in pools did not lead to differentiation of distant present-day European populations, we investigated individual maternal lineages in the following, in order to monitor the connections of the Székely maternal lineages to early Hungarian populations, among others.

On the A12a phylogenetic tree (Figure 5), a modern-day Hungarian sample and a Hungarian sample from the time of the Hungarian conquest (10th century, Harta_HC3), as well as two samples from the 9th–10th century Volga-Ural region (Bolshie Tigani RC8 and Uyelgi-No7, [42,45]), cluster together with the examined modern-day Székely sample. The Conquest Period and the Bolshie Tigani individuals had identical mitochondrial DNA sequences to the Székely individual and that lineage could also be found among today’s Hungarian population.

**Figure 5.**
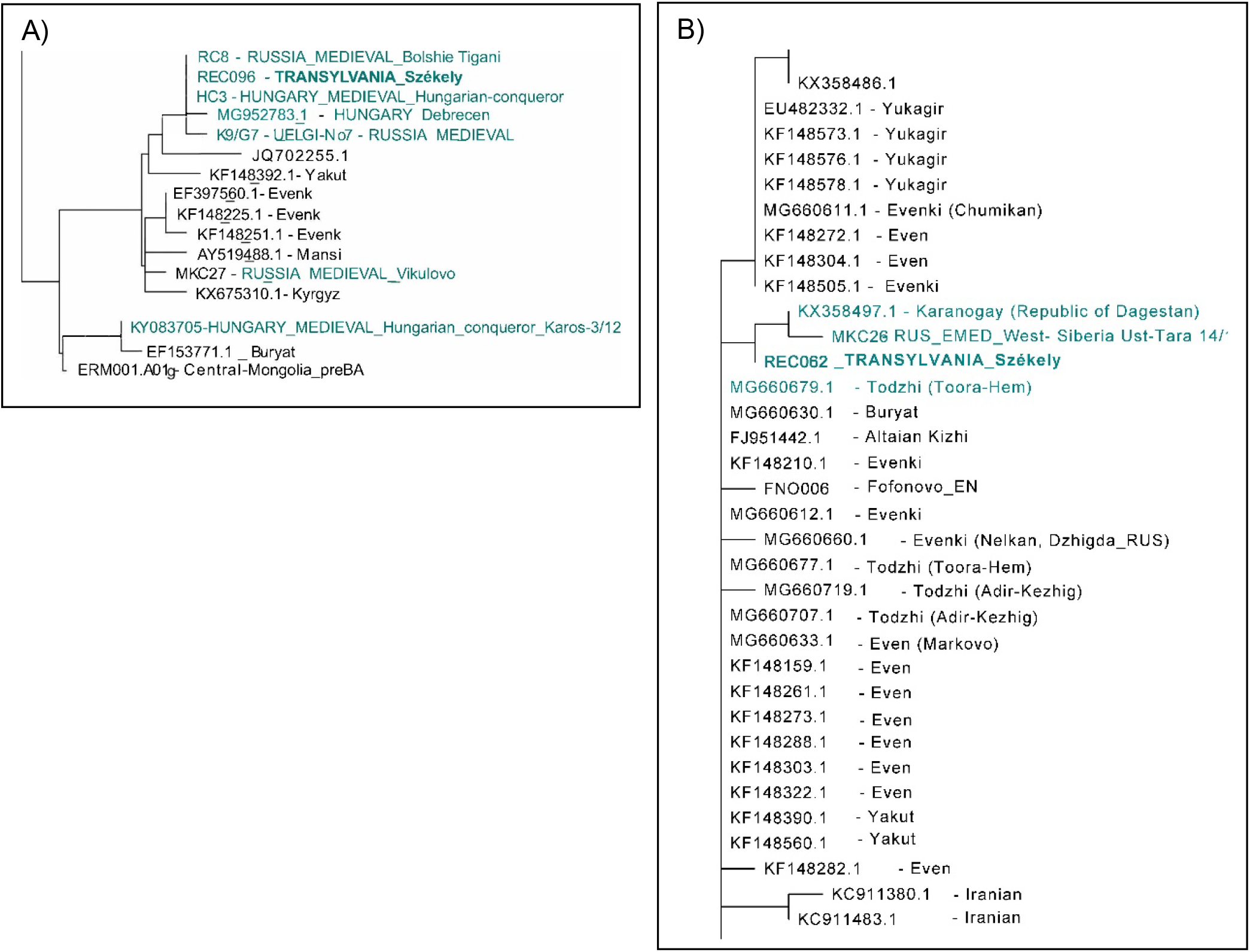
: A) Partial neighbour-joining phylogenetic tree of A12a mitochondrial haplogroup and B) part of the neighbour-joining phylogenetic tree of mtDNA subhaplogroup C4a1a3. Samples highlighted in green are historically relevant to the Székely samples. Most of the data used for the neighbour-joining mitochondrial phylogenetic trees are from the NCBI database; IDs and sources of other data are available in Supplementary Tables S5.

Based on this tree, we assume that the phylogenetic lineage A12a came from the Volga region, and was also present at the time of the Hungarian conquest (late 9th–10th century). The newly reported samples within the A12a subgroup caused some changes in the nomenclature within the A12a tree that we present in the Supplementary Information Figure S4. The Székely sample described here has been ordered to a new subgroup named A12a2b.

On the C4a1a3 partial neighbour-joining phylogenetic tree (Figure 5B) the MKC26 refers to a sample that originated from the 6th–8th century West-Siberian Ust-Tara archaeological site of the Nizhneobskaya culture [45]. The population of this culture was probably proto-Ob-Ugric (Southern-Khanty), although it showed typical Hun-period cultural traits [46]. The other sample which shares a branch with the Székely sample originated from the Karanogay ethnicity (Turkic ethnic group), Dagestan, and the adjacent Todzhi sample was also from a Turkic ethnic group, being a group of Tuvans. This tree represents the mixed nature of the C4a1a3 lineage, which despite its prevalence in Turkic-speaking ethnic groups, may also have originated from Western Siberia in the Székely gene pool. Nevertheless, we do not have immediate proof of that hypothesis in the form of linking lineages from the Volga-Ural region.

Parts of the neighbour-joining phylogenetic tree of A+152+16362 and D4e4 mitochondrial haplogroups are shown in Supplementary Figures S5 and S6. On the A+152+16362 tree, a sample from Cis-Uralic Sukhoy Log cemetery (7th–8th centuries) [45], as well with the latter contemporaneous, ‘Kimak’-reported individual from the Central Steppe [47], can be found in close proximity to the Székely sample. The relationships between samples from the Volga region Early Mediaeval sites Karanayevo, Bolshie Tigani, Gulyukovo, Tankeevka, from the Conquest Period Transdanubia and the Székely sample on the D4e4 phylogenetic tree suggest a possible connection between these sublineages via the conquerors. However, the Székely lineage has a basal character, and is identical to a lineage detected in the Bronze Age of Bolshoy Oleny island in Kola Bay. It is therefore possible that D4e4 is an originally Northeastern European maternal lineage that reached the Carpathian Basin via a different migration.

### Paternal lineages in the new Székely dataset

The population genetic investigation of non-recombining Y-chromosomal markers like Y-STRs and Y-SNPs can be used to trace back paternal lines in time and describe phylogeographic structures and diversities of populations.

#### Haplogroup-based analyses

Y haplotypes from 23 Y-STR markers were obtained from 92 men out of the sampled 115 individuals. The haplogroup predictions were confirmed by selective Y-SNP typing (see methods and Supplementary Table S2). In this dataset, we found eight Y macrohaplogroups (E, G, H, I, J, Q, R and T), which included 21 different subhaplogroups based on SNP typing. Some of these Y subhaplogroups are predominantly found among Inner Asian (R1a-Z93) and South Asian/European Roma (H-M52), as well Northern Eurasian (Q-M242) people (in a total of 7 samples, 7.6% of all samples) the other 18 subhaplotypes are mostly referred to as European-derived types (Table 2).

**Table 2.** Modern-day distribution peaks of the Y-chromosomal subhaplogroups were found in the Székely male population and TMRCA times (given in years before present), based on the Yfull database [50].

| Y chromosome subhaplogroup observed in this study | n | Haplogroup frequency in this study | Modern-day geographic distribution of the haplogroup |
| --- | --- | --- | --- |
| **E-M123** | 2 | 2.17% | Saudi Arabia, Yemen, Europe, Middle East, North Africa - TMRCA 17,700 ybp |
| **E-M78** | 6 | 6.52% | Europe (Mediterranean region), Saudi Arabia, Somalia, Morocco - TMRCA 13,200 ybp |
| **G2-L156 (G2a-P15)** | 4 | 4.35% | Russia, Saudi Arabia, Turkey, Caucasus - TMRCA 18,200 ybp |
| **H-M52** | 1 | 1.09% | India, Pakistan, European Romany populations- TMRCA 15,900 ybp |
| **I1-L22** | 15 | 16.30% | Europe (Sweden, Finland, United Kingdom), North-America - TMRCA 3,900 ybp |
| **I1-M253** | 3 | 3.26% | Europe (Sweden, United Kingdom, Finland, Norway) - TMRCA 4,600 ybp |
| **I2a-P37** | 18 | 19.57% | Eastern Europe (a typical paternal lineage of Mesolithic Europe), Russia - TMRCA 18,400 ybp |
| **I2b-M223** | 2 | 2.17% | Europe (Germany, United Kingdom, Sweden) - TMRCA 14,600 ybp |
| **J2 (M172, L24)** | 2 | 2.17% | Asia, Saudi Arabia, Caucasus, India, Mediterranean, Middle East - TMRCA 27,600 ybp |
| **J2a-M67** | 7 | 7.61% | Russia, Caucasus, Arabian Peninsula, Mediterranean (Italy) - TMRCA 12,100 ybp |
| **J2b-M12** | 3 | 3.26% | Europe (Italy, Germany, United Kingdom), Russia (Volga-Ural region), India, Anatolia, Yemen, Saudi Arabia- TMRCA 15,800 ybp |
| **Q-M242** | 4 | 4.35% | Sweden, Americas, Asia (Russia, China, Mongolia) - TMRCA 28,700 ybp |
| **R1a-M458** | 2 | 2.17% | Poland, Baltic region, Ukraine, China - TMRCA 4,800 ybp |
| **R1a-Z280** | 6 | 6.52% | Asia, (Russia), Poland, Ukraine, Baltic region - TMRCA 4,600 ybp |
| **R1a-Z93** | 2 | 2.17% | Asia (Russia, China), India, Saudi Arabia, Afghanistan, Pakistan, Kyrgyzstan, Turkey - TMRCA 4,600 ybp |
| **R1b-M343\*** | 2 | 2.17% | Europe, North-America - TMRCA 20,400 ybp |
| **R1b-M412** | 1 | 1.09% | Europe (United Kingdom, Ireland, Germany, Portugal, France, Sweden) - TMRCA 5,700 ybp |
| **R1b-P312** | 5 | 5.43% | Europe (United Kingdom, France, Spain, Italy, Sweden) - TMRCA 4500 ybp |
| **R1b-U106** | 5 | 5.43% | Europe (Germany, Sweden, United Kingdom) - TMRCA 4600 ybp |
| **T-M70** | 2 | 2.17% | Syria, Iraq, Saudi Arabia, Mediterranean (Italy) - TMRCA 15900 ybp |

Although some studies on the Székely male populations have been published previously, their comparability with our dataset is rather limited. In 2005, Egyed et al. studied 257 Székely individuals from Miercurea Ciuc, including 89 males typed for 12 Y-STR haplotype loci. In Csányi’s study from 2008, 13 Y haplogroups were determined in the Székely population from Corund [48]. The Y haplogroup diversity was 0.9157 in the latter Székely population, 0.9011 in the Székelys living in Miercurea Ciuc [7] and 0.8636 among the Hungarian male population in Hungary [14]. The Y-STR haplotype diversity in our studied Székely population was 0.9995, and the proportion of unique haplotypes was 97.8% using the PowerPlex Y23 System. Based on an investigation of 72 European populations comprising a total of 12,000 samples, the average haplotype diversity was higher (Hd = 0.999992) using the same Y23 System [49] than in the Székelys.

In 2015, Bíró et al. studied Székely haplogroups from Miercurea Ciuc (haplotypes published by Egyed et al.) [9,11] that had proven Central and Inner Asian genetic contributions: in their dataset, the possible maximum Central/Inner Asian admixture among the Székely male population was 7.4% [9,11]. In our results, this proportion was a comparable 7.6% in the population around Odorheiu Secuiesc. According to Bíró’s study of contemporary Hungarians from Hungary, this Central/Inner Asian admixture was estimated as only 5.1%, and 6.3% in the Csángó male population from Ghimeş. Examining the Bodrogköz Hungarian dataset, these Asian-derived haplogroups appeared in 6.9% of the population.

The comparative analyses of the Y haplogroups with other Székely populations showed some level of diversity among the Székely groups. However, while N1a [42,45,51,52] occurred among the Székelys in Miercurea Ciuc, the population of the Odorheiu Secuiesc region did not yield such a signal in our analysis, but resulted in higher proportions of haplogroups Q, I and I2a than in the previously studied Székelys (Figure 6).

**Figure 6.**
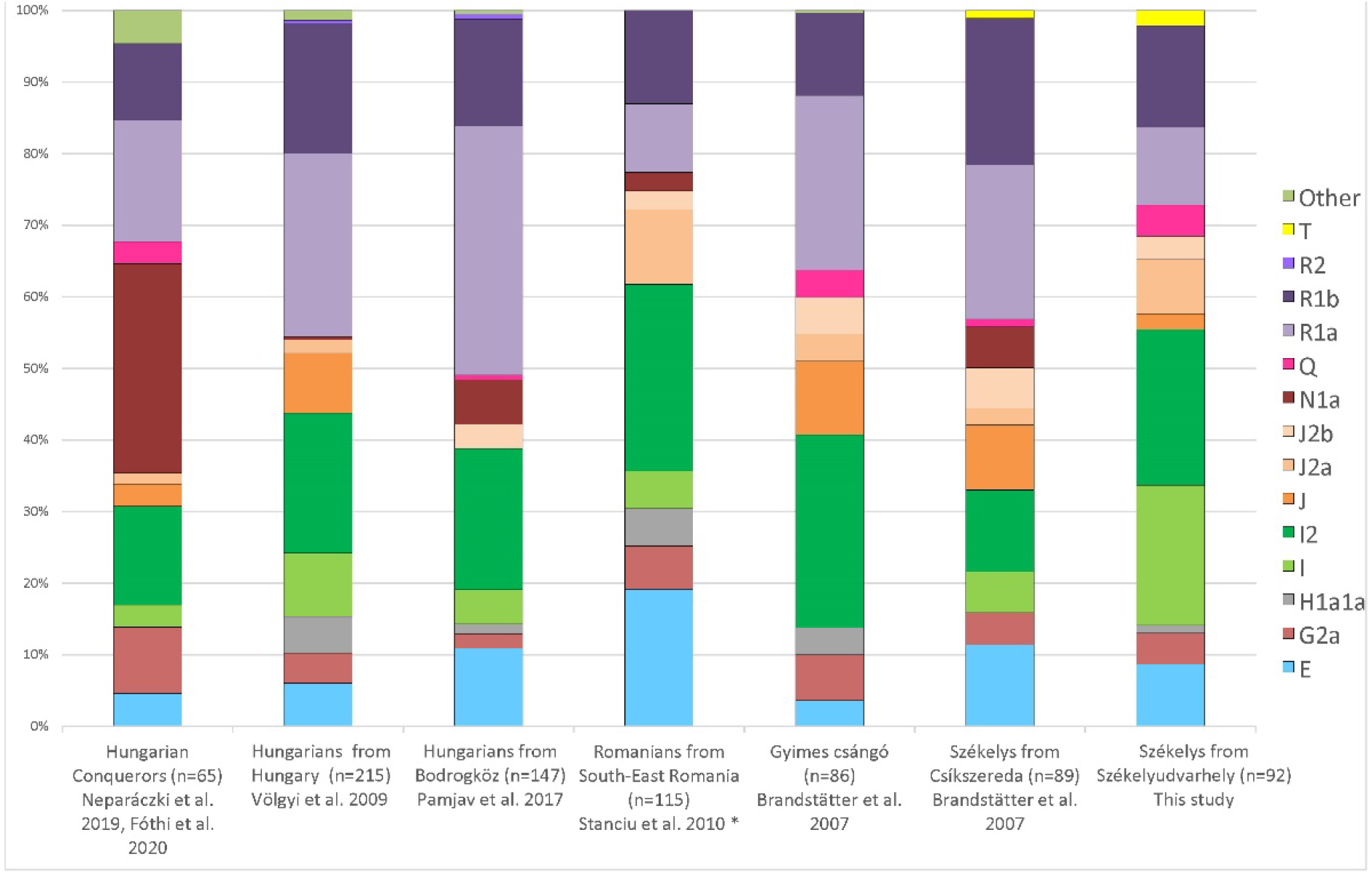
Diagram of the Y haplogroups in Székely, Ghimeş Csángó, Hungarian and Romanian populations. *Haplogroups from the Romanian population were predicted from 17 STR data using nevgen.org.

The haplogroup R1a comprised a substantially higher proportion of the haplogroups among the Hungarian populations living in Hungary than among either the Székely population of the present study, or the Ghimeş Csángó and Romanian populations. In the Székely population we studied, R1a and R1b comprised only 25% of all haplogroups, while in the case of Hungarians in Hungary, they accounted for 45–50%. However, the ratio of I and J2 haplogroups was higher in the studied Székelys than in the Hungarians. Furthermore, the frequency of haplogroup J was higher in the Székely population of Miercurea Ciuc than in Odorheiu Secuiesc.

Székelys in Miercurea Ciuc showed roughly similar frequencies of G2a and E haplogroups to Székelys around Odorheiu Secuiesc; furthermore, the T haplogroup did not appear elsewhere in the comparison except for in the two Székely groups. T-M70 seems to have originated from the Fertile Crescent and possibly arrived in Europe in the Neolithic with the first farmers [53,54]; today it shows the highest frequency in East Africa and the Middle East. The Székely T-M70 samples belonged to the T1a2b1 subhaplogroup, which is rarely found in ancient data; this was made even more interesting given that it was found in the Hungarian Conquest Period horizon of the Western Hungarian Vörs-Papkert cemetery [55,56].

The I2a1a (I2a-P37) haplotype occurred in the highest number in the studied Székelys, comprising 20% of the total haplotypes. In Hungary, 16.74% of men carry this haplotype [14]. I2a-P37 and subgroups occur at high frequencies in Sardinia (38.9% [57]), and are also present at high frequency among Balkan populations [58]. The proportion of the I2a1a Y type in the Romanian population is 17.7%.

Another dominant haplogroup in our dataset is I1a1b1 (I1-L22), which is most frequent in Sweden and Finland, and represents a fairly large Nordic branch of I1. It was dispersed by the Vikings and nowadays can be frequently found in the Baltics, Britain, Poland and Russia [59].

The distribution of Y haplogroups between the villages did not show a characteristic pattern or patrilineal system; the observed haplogroups were mixed within the region, as is demonstrated in Supplementary Figure S7.

#### Y chromosome phylogenetic analyses

Our dataset contained four samples that belonged to the Y-chromosomal haplogroup G2a, which we analysed in further detail. In the absence of comparative data covering 23 STR, the network in Figure 7 is based on 17 STR. Two of the Székely G2a Y chromosomes clustered with the M406 subgroup of G2a, with individuals sampled in Tyrol, Austria [60]. This terminal SNP defines the Y-chromosomal subgroup G2a2b1 (ISOGG 2020 v15.73) [27], whereas some of the Székely samples most probably belong to G2a2b2a1a1b based on the L497 marker (ISOGG 2020 v15.73). The G-L497 sub-haplogroup likely originated from Central Europe and has been mostly prevalent in European populations since the Neolithic period [54][61]. The G-L497 lineage could potentially be associated with the *Linearbandkeramik* (LBK) culture of Central Europe. The G-M406 sub-cluster is most concentrated in Cappadocia and Anatolia in Turkey nowadays [62], and has been present in that area since the early Neolithic [63,64].

**Figure 7.**
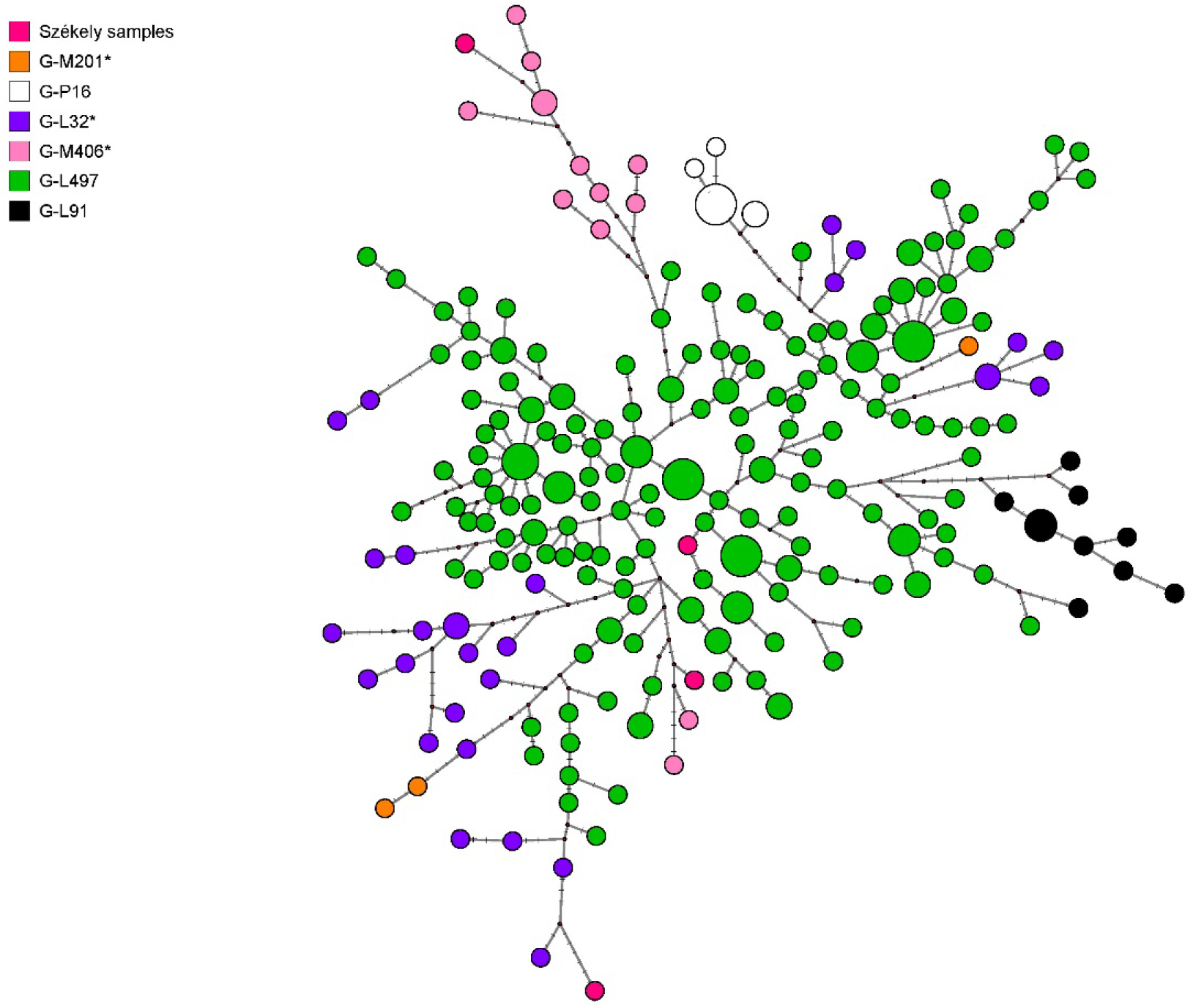
G2a median-joining network: The network was constructed based on 17 Y-STRs, the data used for the network are all from the Tyrolean region of Austria [60]. Circles represent distinct haplotypes, the size of the circles is proportional to the haplotype frequency (the smallest circle corresponds to one individual), the colouring corresponds to haplogroup subclades where the Székelys are highlighted.

Our dataset comprised six samples classified into the Y chromosome R1a1a1b1a2 (R1a-Z280) haplogroup, which we visualised on a median-joining network (Figure 8). For comparison, we collected Y-STR data from the Family Tree Y-DNA database R1a page and we filtered the samples for Y4459 SNP based on nevgen.org prediction. In addition, we included data with the same haplogroup classification from Hungary, Bodrogköz [15] and three early mediaeval samples from the Volga-Ural region (Novo Hozyatovo, Gulyukovo—Chiyalik culture), which may have incorporated Hungarians who remained in the East after the Hungarians Westward migration [65][45].

**Figure 8.**
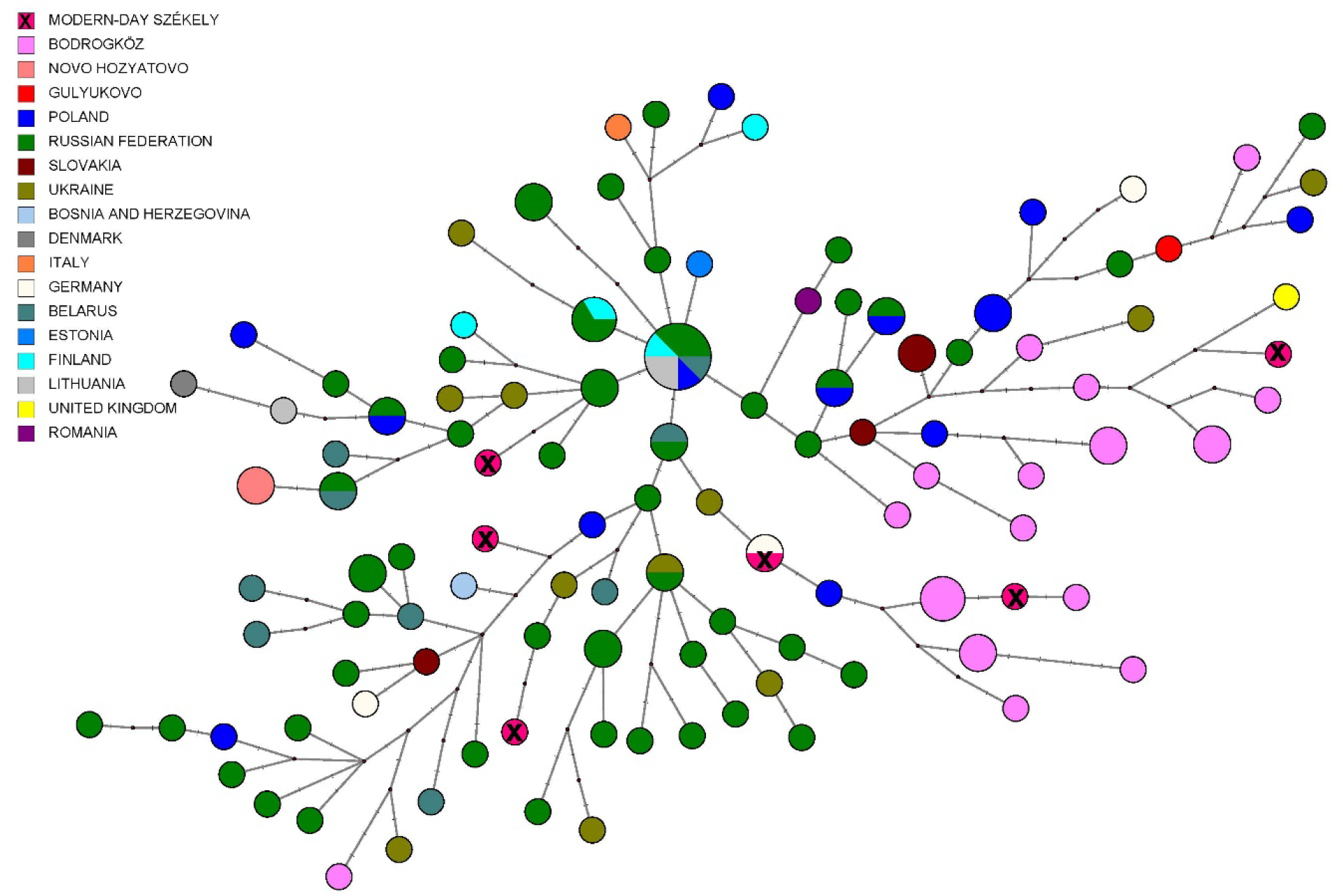
R1a-Y4459 median-joining network. The analysis was performed based on 15 STRs. No larger clusters are seen on the figure, but close relationships can be detected between the samples. Near the Székely samples, individuals from the Bodrogköz region, Russia, Germany and Poland can be found.

We present further median-joining networks in the Supplementary Data, Figures S8-S10, for haplogroups Q-M242, R1a1a1b1a1a (R1a-M458) and R1a1a1b2 (R1a-Z93). The network of Q-M242 placed the present Székely samples among the Bukovina-Székelys, whereas the two R1a networks did not show geographically relevant patterns.

There were four samples in our dataset that could be classified into the Y-chromosomal macrohaplogroup Q, three of which belonged to the Q1a-F1096—probably to subhaplogroups Q1a2-M25 and Q1a2a1-L715—and one which belonged to the subgroup Q1b1a3-L330; these subgroups are interesting, from our perspective, due to their Central Asian origin (Supplementary Figure S8). Q1a2-M25 is known from the second half of the 5th century AD near Sângeorgiu de Mureş, Romania [66]. Based on the discovered grave goods of the buried man at this site, and his Asian cranial features as well as artificial cranial deformation, strong Hun period traditions have been pointed out [66]. The Q1a2-M25 lineage was also demonstrably present in the Carpathian Basin in the first half of the 7th century AD from a richly furnished, high-status Avar horseman warrior’s grave in the Transztisza region, belonging to subhaplogroup Q1a2-M25 [67,68]. Ancient individuals with the same Y-chromosomal haplogroup are known from the Early Middle Bronze Age Okunevo and from the Baikal Early Bronze Age (Shamanka and Ust’Ida sites), and the Tian Shan Hunnic [47] and Hungarian Sarmatian cultural context [68] as well.

The Q1b1a3-L330 subhaplogroup was also present from the middle third of the 7th century AD in the Carpathian Basin; it was identified from a richly furnished early Avar grave [67,68], and according to a median-joining network (Figure S5 in [67]), this male had a probable Altaian or South Siberian (Tuvinian) paternal genetic origin. The Q1b1a3-L330 lineage was present in the Proto-Ob-Ugric group (Ust’-Ishim culture—Ivanov Mis and Panovo sites), which corresponds to its Altai or Siberian origin [45,69]. The genetic imprint of the Avars in the Székely population can have multiple origins, as their 8th-century settlements were scattered throughout the central Carpathian Basin and along the Maros River in Transylvania, and part of them probably persisted after the Avar–Frankish wars as well [1].

The R1a-M458 median-joining tree shows notable separation of the R1a-M458 types within the Hungarian Bodrogköz population, and the connection of the Székely lineages to some parts of it. On the R1a-Z93 median-joining tree Hungarian King Béla III and other skeletal remains originating from the Royal Basilica of Székesfehérvár [70,71] show a great genetic distance from the Székely samples, just like the Bashkirian Mari males (Figure S10).

Examining the R_ST_ values based on Y-STR profiles (see Supplementary Table S6, Supplementary Information Figure S11), the closest population to the Székelys was the Slovenian group, with non-significant R_ST_ *p*-value; the populations with significant R_ST_ *p*-values were the Greek, Hungarian, Hungarians from Bodrogköz, Serbian and Croatian datasets (Figure 9). All these reflected a strongly Southeastern European-funded base population of the Székelys with a limited proportion of surviving eastern elements.

**Figure 9.**
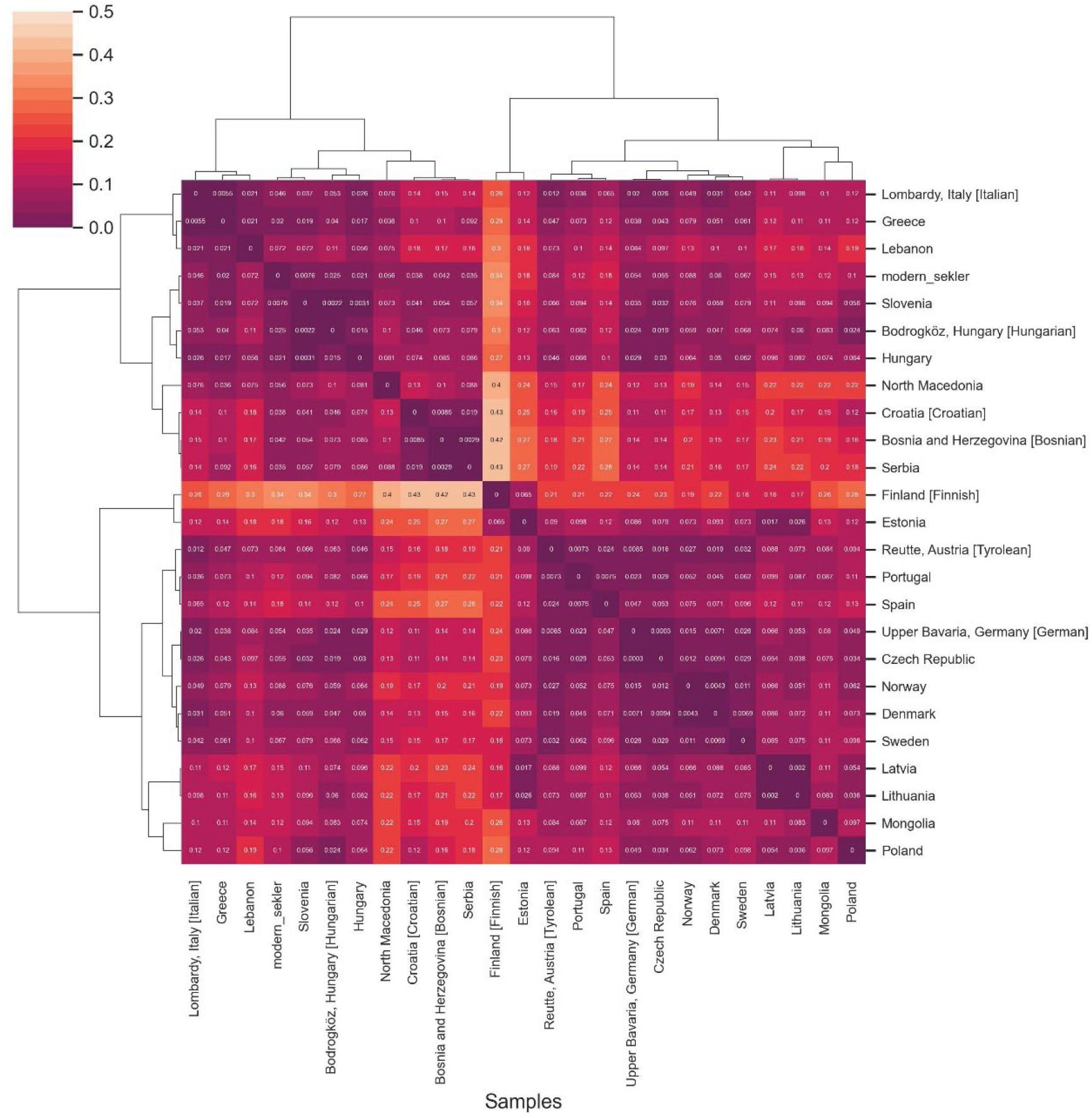
Heatmap of pairwise R_ST_ values with clustering applied for the modern Székely group and populations from Europe (colour scale ranging from yellow to dark purple). We calculated it in Python using the seaborn clustermap function with parameters ‘correlation’ distance metric and ‘complete linkage’ method [35].

#### Comparison of paternal lines with ancient data

Most of the paternal lines of the early mediaeval sites in the Western-Siberian and Volga-Kama regions—which are suspectedly linked to Hungarian ethnogenesis—belong to N1a1-M46, Q1b-M346-L330, G2a2b-M406, R1a and J2a1-Z6046 [45].

According to the publications of Fóthi et al. [52], Neparáczki et al. [51] and Csányi et al. [48], the N1a1-M46, R1b-U106 and I2a-M170 lines were the most widespread among the conquering Hungarians they examined. This suggests that the conquerors were of diverse origins and, while N1a1-M46 originated from the east, the R1b-U106 (R1b1a1b1a1a1) lineage is known from the Late Copper Age/Early Bronze Age transition in Europe [72] and is most prevalent in Germanic-speaking people nowadays [73].

These observations fit to the genomic data published by Maróti et al. [56]; among the group they define as the ‘conquering elite’, the haplogroups N1a1a1a1a2a1c (Y13850), N1a1a1a1a4 (M2128), I2a1a2b1a1a (YP189) and R1a1a1b2a (Z94) can be found in notable frequencies.

According to all previous studies, the N1a1 line is the most characteristic of the Hungarian conquerors—almost 30% of the lines belong to the N1a group, and also appear with lower frequencies (4–6%) in the Bodrogköz and Miercurea Ciuc populations; however, this line is completely missing from our new Odorheiu Secuiesc region dataset. The second most widespread haplogroup among the Hungarian conquerors is the R1a-M198 (16.9% of all haplogroups) which is quite common nowadays in the Odorheiu Secuiesc region (10.9%). R1b and I2 haplogroups are present in a relatively higher proportion both in the conqueror (10.8% and 13.8%, respectively) and Székely groups (14.1% and 21.7%, respectively). Most of the other haplogroups listed above—like R1b-U106, G2a and Q1b-M346-L330—can also be found among the Székelys, at a maximum frequency of 5%. The direct comparison of the data is limited, however, by the numerous allelic dropouts in the ancient STR analysis [52], and by the different levels of haplogroup resolutions obtained.

## Discussion

In this study, we presented the maternal and paternal genetic composition of a Székely group, a Hungarian-speaking minority living around the city of Odorheiu Secuiesc in Transylvania (Romania). We carefully selected 115 sample providers with local ancestors inhabiting small villages in the area. Altogether, 115 complete, high-coverage mitochondrial genomes were produced with next-generation sequencing methods, which revealed 89 unique haplotypes and could be classified into 72 different subhaplogroups. These mitochondrial haplogroups are mainly present in European regions, but there are also some Asian- and Near Eastern-derived lineages, like A+152+16362, A12a, C4a1a3, C5c1a and D4e4.

In this new Székely dataset, the discovery of an Asian maternal lineage (A12a) completely identical to that found in a male with typical Hungarian conqueror artefacts from the 10th-century cemetery in Harta, Hungary [74], and in the early Hungarian cemetery of Bolshie Tygani in the Volga-region, is a robust sign that some lineages in the Székely population are shared with Hungarian conquerors and are thus most probably of common origin.

The 92 paternal lines investigated in the dataset were mainly composed of European haplogroups, but some lineages (I1-L22, T-M70, J2a-M67) stood out or showed a different distribution in their proportions than in other surrounding Székely ethnic groups. The performed Y-STR networks allowed detailed observations on paternal lineages G2a-L156, R1a-Z280, Q-M242, R1a-M458 and R1a-Z93.

We detected a strongly Southeastern European base population of the Székelys. The genetic proximity of Balkan populations may be a consequence of the formerly inhabited areas of the Székelys in southern Hungary. Presumably, the signs of Glagolitic and Cyrillic origin included in their own old script are also connected with the Eastern European and Balkan connections of the Székelys. This alphabet was probably developed in the Carpathian Basin during the 10th century—using certain signs of the Turkish runic script too—and became suitable for writing short texts in Hungarian. It was used exclusively by the Székelys [75].

The Hunnic origin of the Székelys remains questionable in the light of the present genetic data, because we have scarce genetic data from the Hunnic period [51,68]; furthermore, the population living in the Carpathian Basin during the Hunnic period and in the Avar period (before the time of the Hungarian conquest) shows large heterogeneity [68]. Here we demonstrated connections of the Székelys to the 5^th^-7^th^ centuries’ population through Y-chromosomal Q Asian lineages, which, however, could have arrived repeatedly in the region in numerous epochs. The possible separation of the different immigrant waves requires larger comparative databases from the early mediaeval and mediaeval periods.

We found large among-village heterogeneity in both the maternal and paternal gene pool. The current Székely dataset completes the previous studies and is broadly in line with their observations. The genetic connections between the Székelys and Hungarians could be detected based on our uniparental genetic data using allele frequency analyses; moreover, genome-wide, haplotype-based methods [12] confirm our observations. The results of this genome-wide, autosomal marker-based analysis show that Székelys from Corund and non-Székely Hungarians from Transylvania are strongly related to contemporary Hungarians living in Hungary [12].

Compared to previous uniparental studies involving Székelys [6–9,11,48], what is different and novel in our research is the sampling method, the selection of the participants and the careful documentation of the ancestors. The results of our research and the reported data are definitely a qualitative leap, considering that so far complete mitochondrial DNA data have not been available from the region and Y-chromosomal data containing 23 STRs have not been previously reported.

Besides revealing present-day diversity, it is of great importance to evaluate genetic continuity or transformation between present-day and ancient populations. To explore this, further mediaeval samples, regional genetic transects and complete genome analyses are aimed for. The follow-up project involves the study of mediaeval cemeteries from the same Odorheiu Secuiesc region to monitor the population history of the Székelys.

## Author Contributions

Conceptualisation, A.Sz-N. and B.G.M. and E. B and I.M.; Methodology, O.Sz. and N.B and B.Sz and D.G.; Formal Analysis, Writing – Original Draft Preparation, N.B. and O.Sz.; Writing – Review & Editing, E.B. and H.P and N.B. and A.Sz-N.; Visualisation, N.B. and O.Sz and A.Sz-N.; Supervision, A.Sz-N. and H.P and B.E.; Funding Acquisition, A.Sz-N.

## Institutional Review Board Statement

For sampling, handling and storage of personal data and genetic samples, we considered the Hungarian 2008/XXI. law as guidelines. The Hungarian 2011/CXII. law provided us with rules about the information and self-determination rights of the sample providers. The research plan was forwarded to the Deputy State Secretary for National Medical Officers in the Ministry of Human Capacities, and to the Committee on Research Ethics for their consent.

## Informed Consent Statement

Informed consent was obtained from all subjects involved in the study.

## Acknowledgement

The authors would like to thank all voluntary sample donors and the organisers of the sampling for their contribution to the project.

## Author Approvals

All authors have seen and approved the manuscript, and it has not been accepted or published elsewhere.

## Abbreviations

AMOVA: Analysis of Molecular Variance
HG: Haplogroup
MDS: Multidimensional Scaling
MJ: Median-Joining
NGS: Next-generation Sequencing
NJ: Neighbour-Joining
PCA: Principal Component Analysis
SNP: Single Nucleotide Polymorphism
STR: Short Tandem Repeat
Y-STR: Y chromosome Short Tandem Repeat
YHRD: Y-STR Haplotype Reference Database
Ybp: years before present

