## supplementaryfigures for "High Coverage Mitogenomes and Y-Chromosomal Typing Reveal Ancient Lineages in the Modern-day Székely Population in Romania"

**Supplementary Information**

Noémi Borbély, Orsolya Székely, Bea Szeifert, Dániel Gerber, István Máthé, Elek Benkő, Balázs Gusztáv Mende, Balázs Egyed, Horolma Pamjav, Anna Szécsényi-Nagy

Figure S1: Ward Hierarchical Clustering

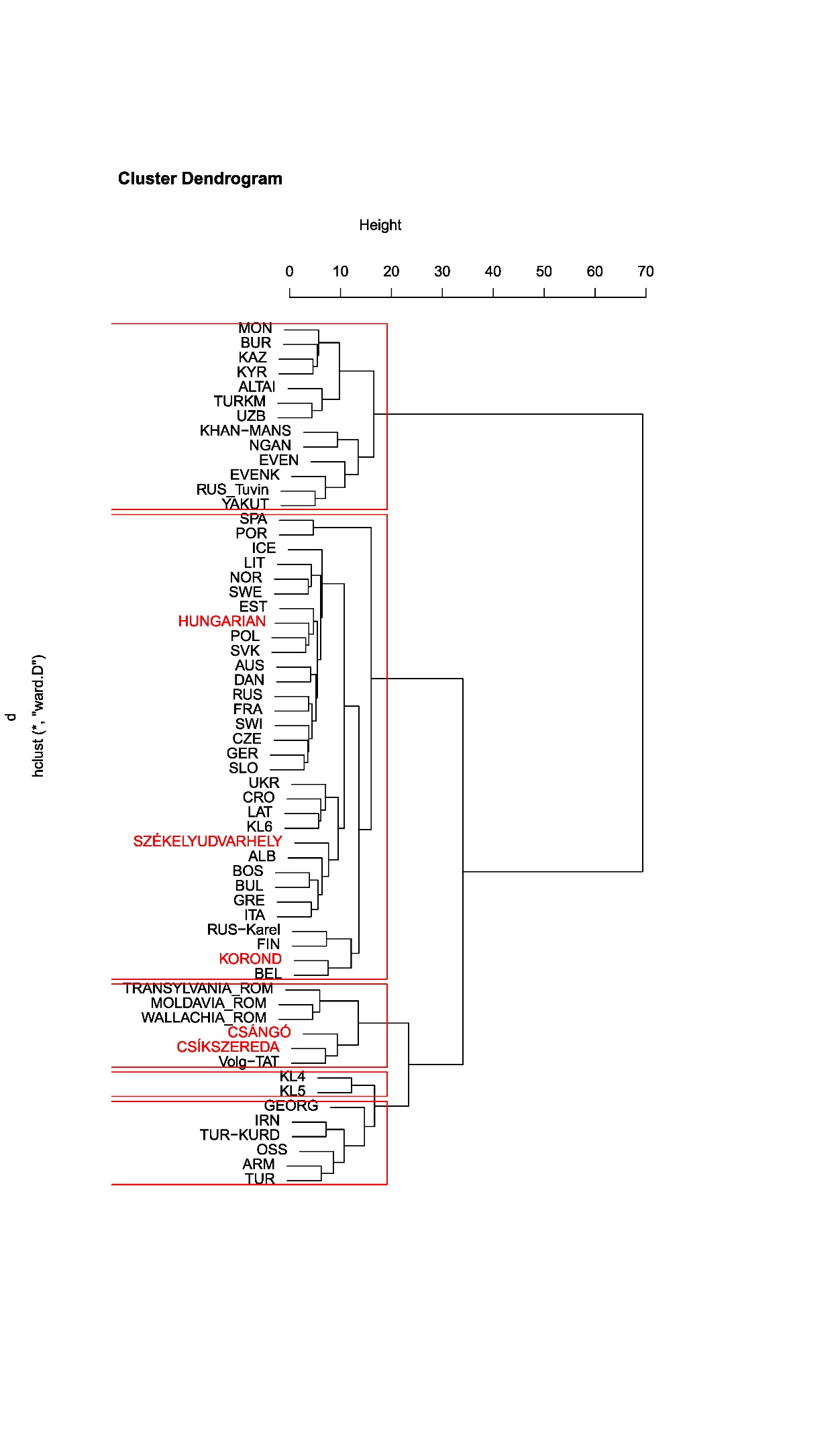

Figure S1: Ward Hierarchical Clustering. The result of the Ward cluster analysis is based on PC1-PC6 scores calculated for haplogroup frequencies of 56 modern-day and three ancient populations’ (the datasets are the same as used for the PCA analysis). The results show that the investigated Székely group forms a sub branch with modern European populations: Albanian, Bosnian, Bulgarian, Greek and Italian groups are on the same branch.

Figure S2. PCA plot with 56 modern and three ancient populations

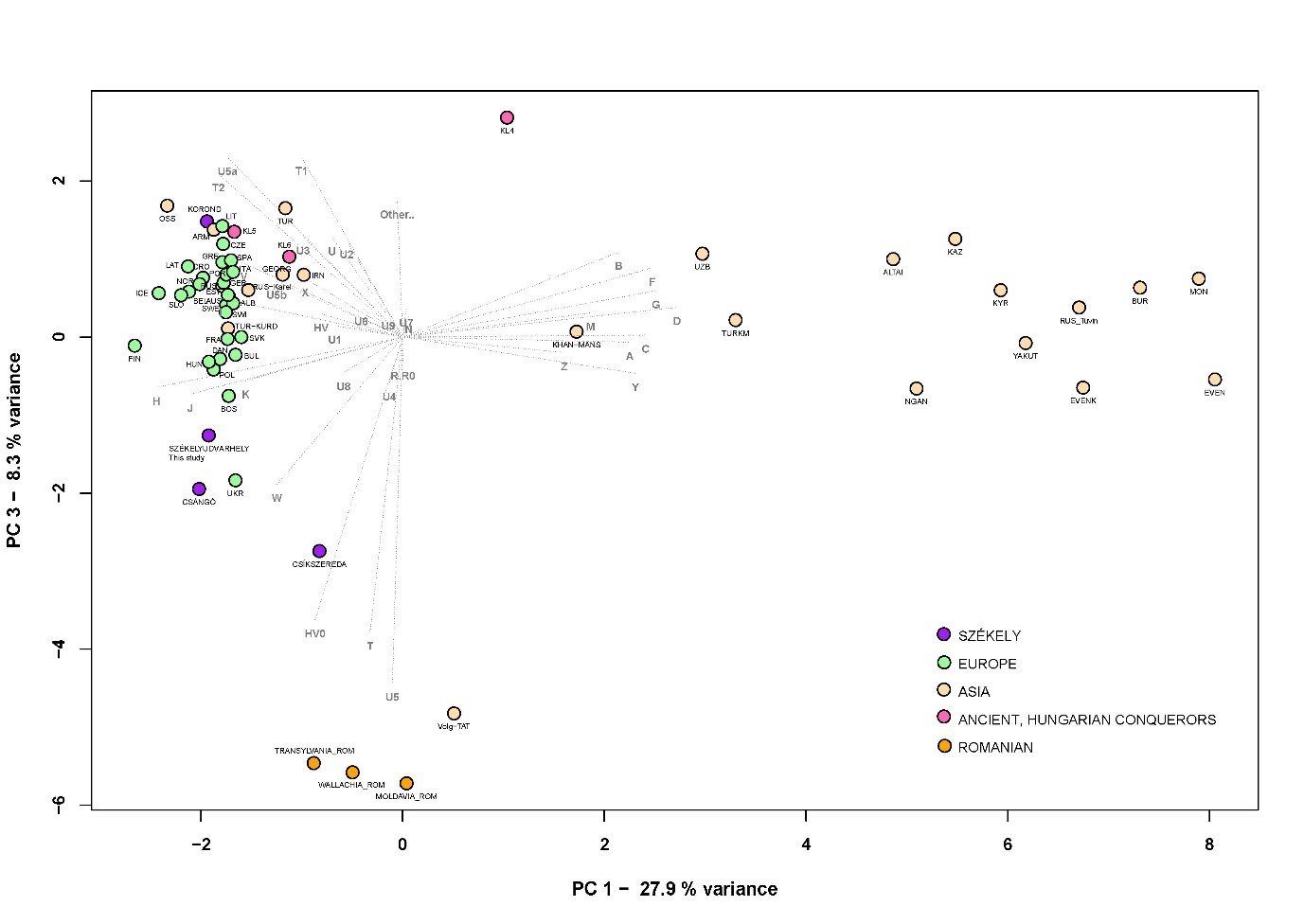

Figure S2. PCA plot with 56 modern and three ancient populations, representing ﬁrst and third principal components (36.2% of the variance). PCA analysis based on haplogroup frequencies in Eurasian modern populations and three ancient populations (Hungarian conquerors, group KL4-6 based on Kovács, 2013 [1], see Supplementary Table S3) from Hungary. The investigated Székely population and previously examined Székely groups are marked in purple, the ancient populations from Hungary are indicated in pink, Romanian populations are in orange. Europeans are colored green, Asian populations have a drab colour. The Székely populations which clustered together on the PC1-PC2 plot split up on the PC1-3 plot while the Romanian groups located further to the Székely groups along the PC3 axis.

Figure S3. Whole mitogenome sequence-based evaluation: heatmap of pairwise F_ST_ values for the modern Székely group and 37 reference populations

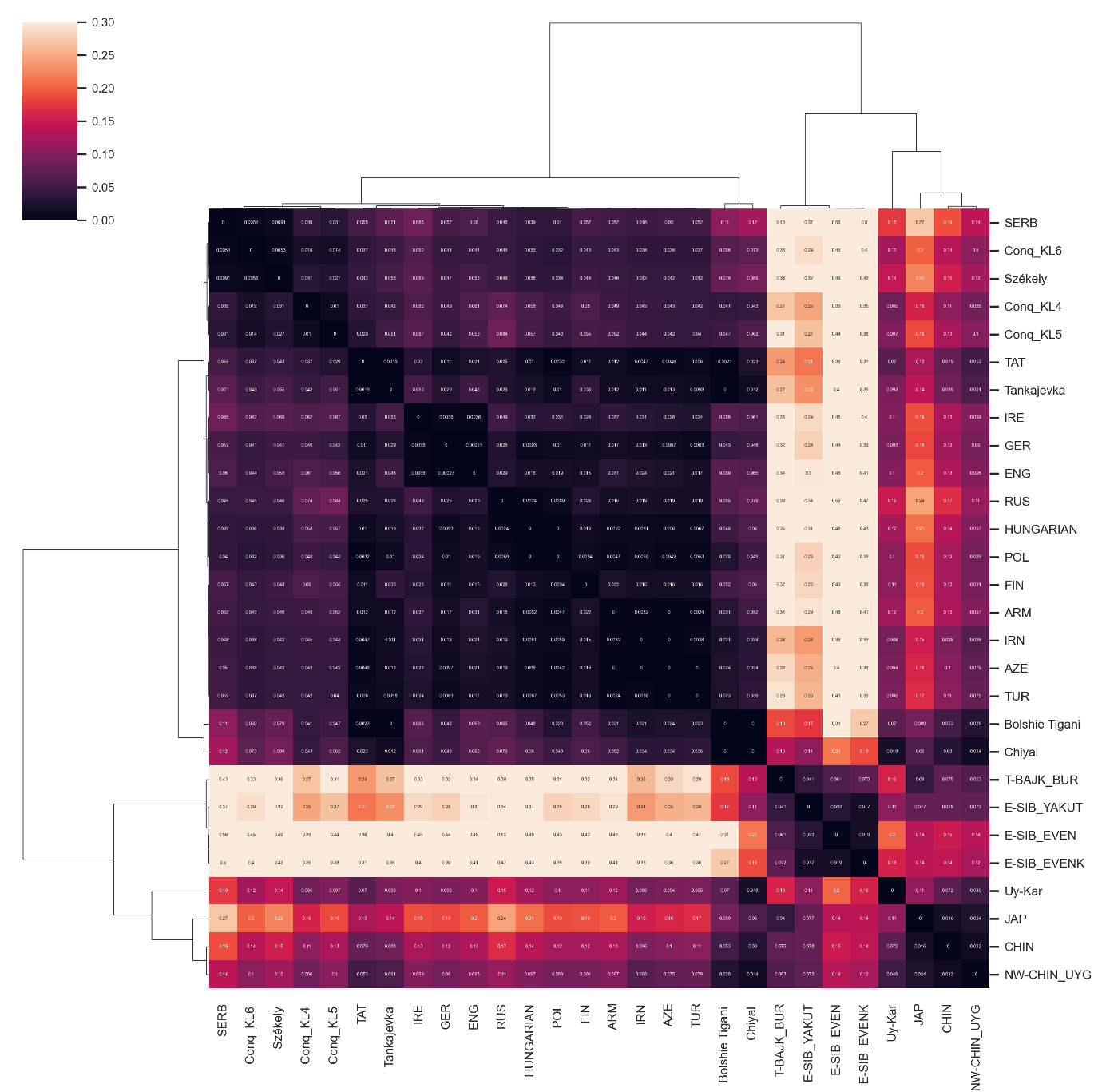

Figure S3. Heatmap of pairwise F_ST_ values for the modern Székely group and 27 reference populations with a colour scale ranging from yellow to dark purple. The lighter block colours indicate larger genetic differentiation, whereas the darker colours show closer genetic affinities between the pairs of populations. The European groups all show great similarities to each other. The Székelys cluster on the European branch with Serbians and conqueror period Hungarian groups; in addition to these, the Hungarian and Polish groups show the closest links. We calculated the clustermap in Python using the seaborn clustermap function with parameters: metric = ‘correlation’, method = ‘complete’.

KL6 refers to cemetery group 6 based on Kovács, which represents cemeteries of large villages opened in the 10th century and used until the 11th and 12th centuries [1]

Figure S4. Phylogenetic tree of mitochondrial group A12a.

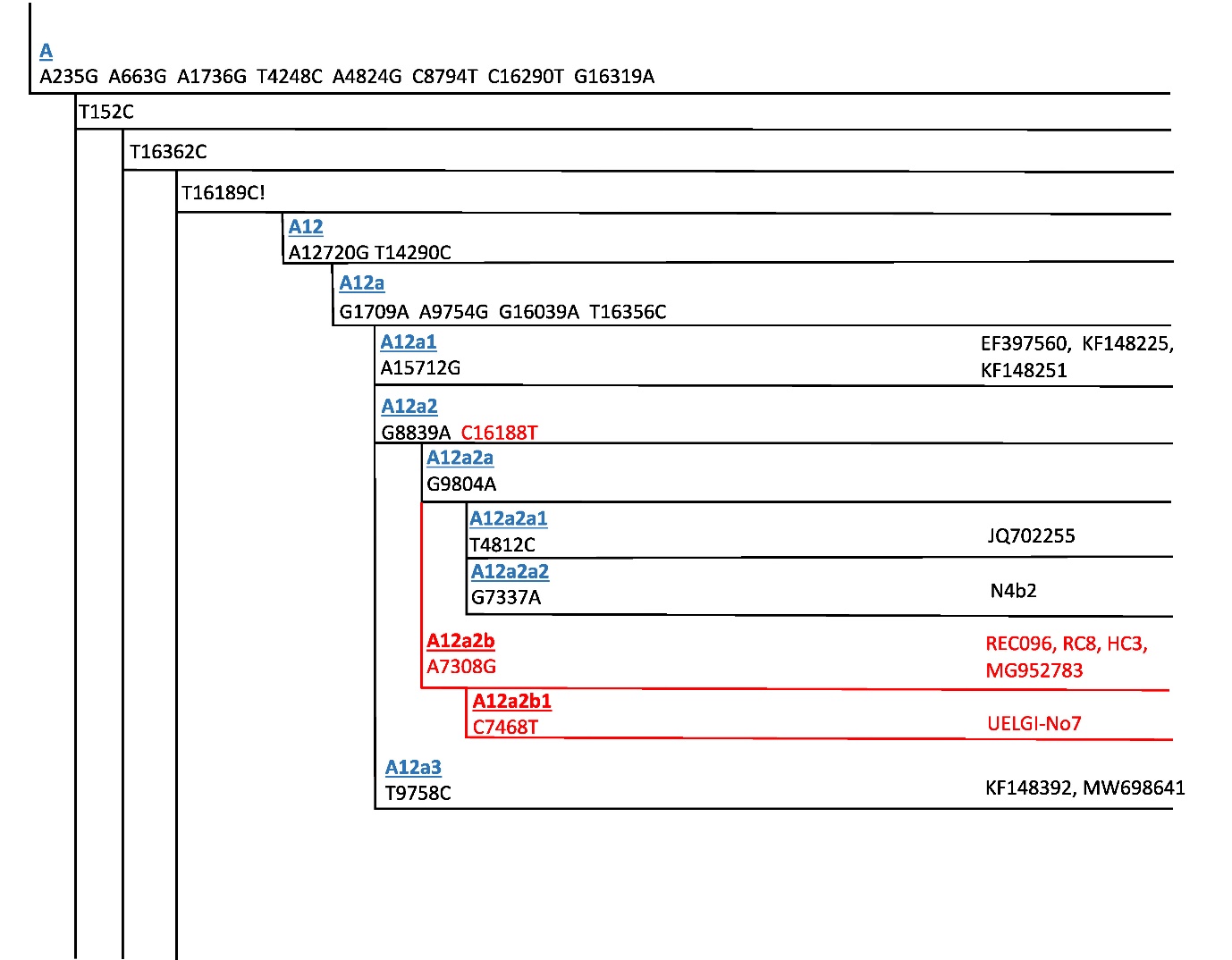

Figure S4. Phylogenetic tree of mitochondrial group A12a. Taking into consideration the currently used nomenclature of the phylogenetic tree of global human mitochondrial DNA variation we described new subbranches on the A12a tree, named as A12a2b and A12a2b1. These fit perfectly into the current nomenclature, the new subbranches do not affect the previously named samples and branches. All data were used from [MTree](https://www.yfull.com/mtree/A12/) [2], and Ian Logan mtDNA [3], the codes of the samples are indicated at the end of each line. The new branches are coloured in red, the A12a2b group is made up of samples from the investigated Székely group (REC096, modern-day sample) from Hungary (MG952783: Hungarian present-day sample; HC3: mediaeval, classical Hungarian conqueror sample), and from Bolshie Tigani and Uyelgi (RC8 [4], and UELGI-No7 (see in Csáky et al, 2020 [5], under ID SB7) mediaeval Hungarian-related individuals).

Figure S5. Part of the neighbour-joining phylogenetic tree of mtDNA subhaplogroup A+152+16362

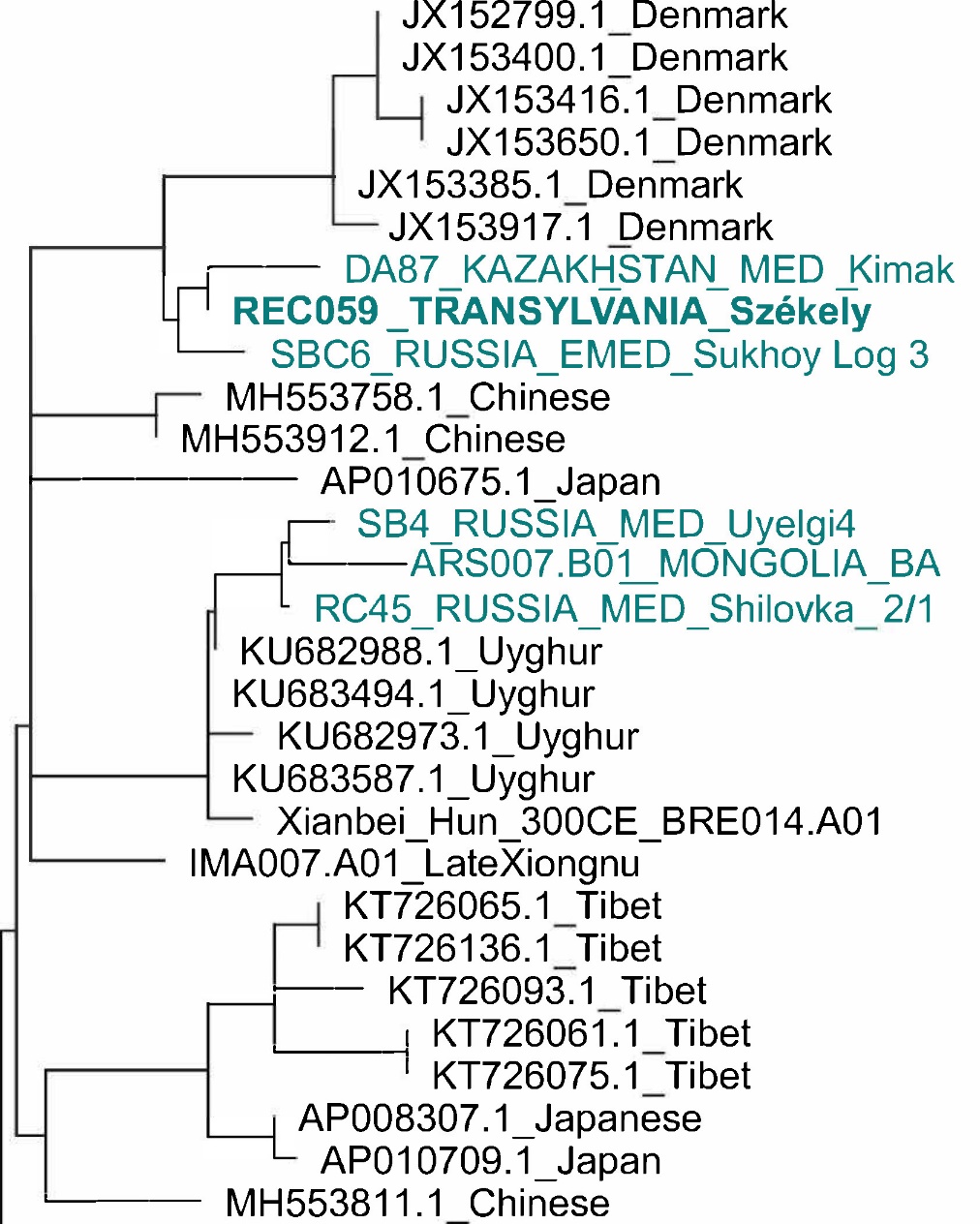

Figure S5. Part of the neighbour-joining phylogenetic tree of mtDNA subhaplogroup A+152+16362. On the A+152+16362 phylogenetic tree a sample from 7–8th centuries Sukhoy Log cemetery from the Cis-Ural (SBC6_RUSSIA_EMED_Sukhoy Log 3), as well as a mediaeval Kimak culture related individual from Central Steppe (KAZAKHSTAN_MED_Kimak_DA87) can be found, which are in close proximity to the Székely sample (REC59_TRANSYLVANIA_Székely). On a branch near the REC059, many Uyghurs are located, and one mediaeval sample is from the eastern side of the Urals (RUSSIA_MED_Uyelgi4), another mediaeval sample from the Volga region Novinki group, Shilovka site (RUSSIA_MED_Shilovka_RC45) and one from the Bronze Age of Mongolia. These connections can be traced back to the eastern steppe environment and suggest a Central-Eastern Asian origin of this maternal lineage.

Figure S6: Partial D4e4 neighbour-joining phylogenetic tree

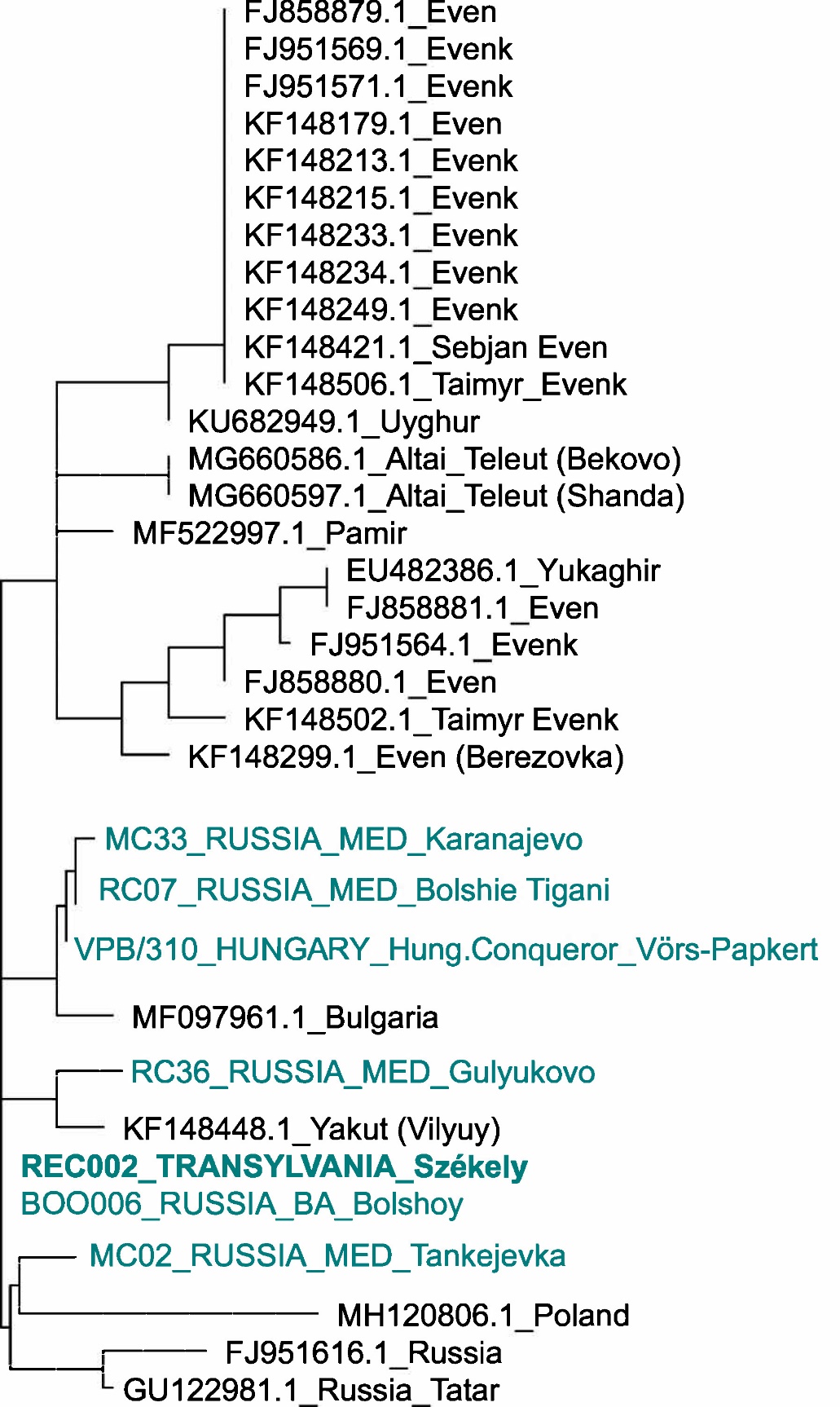

Figure S6: Partial D4e4 neighbour-joining phylogenetic tree. The relationships between lineages from Karanajevo, Bolshie Tigani, Gulyukovo, Tankejevka, Western Hungary from the Conquest Period, and a Székely on the phylogenetic tree suggest a pre-conquest period connection of these lineages.

Figure S7. Y-network based on 23 STR data from the modern Székely population

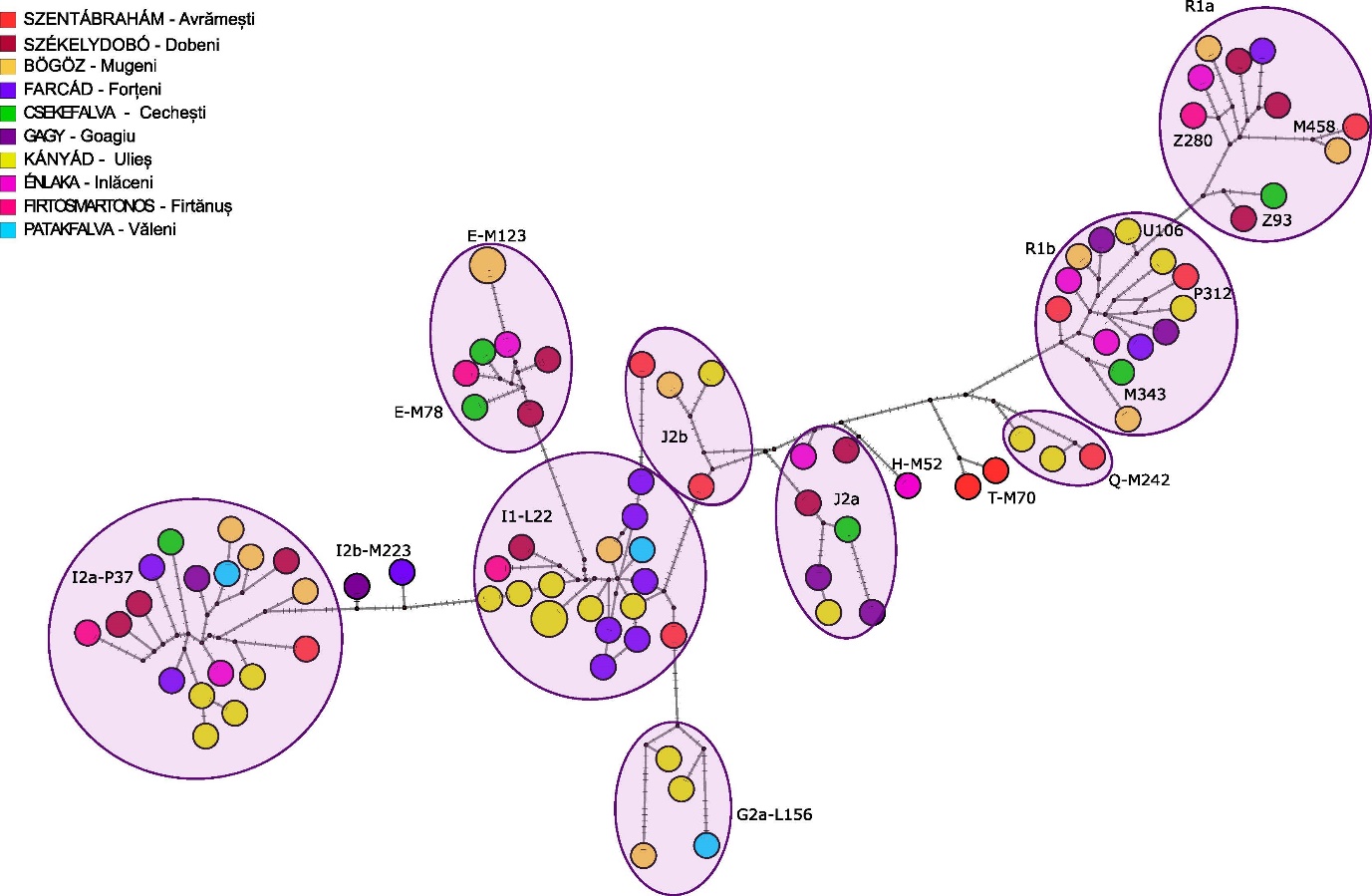

Figure S7. Y-network based on 23 STR data from the modern Székely population (n=90). The colours indicate the villages where the sample providers live at the time of sampling.

Figure S8: Q- M242 median-joining network

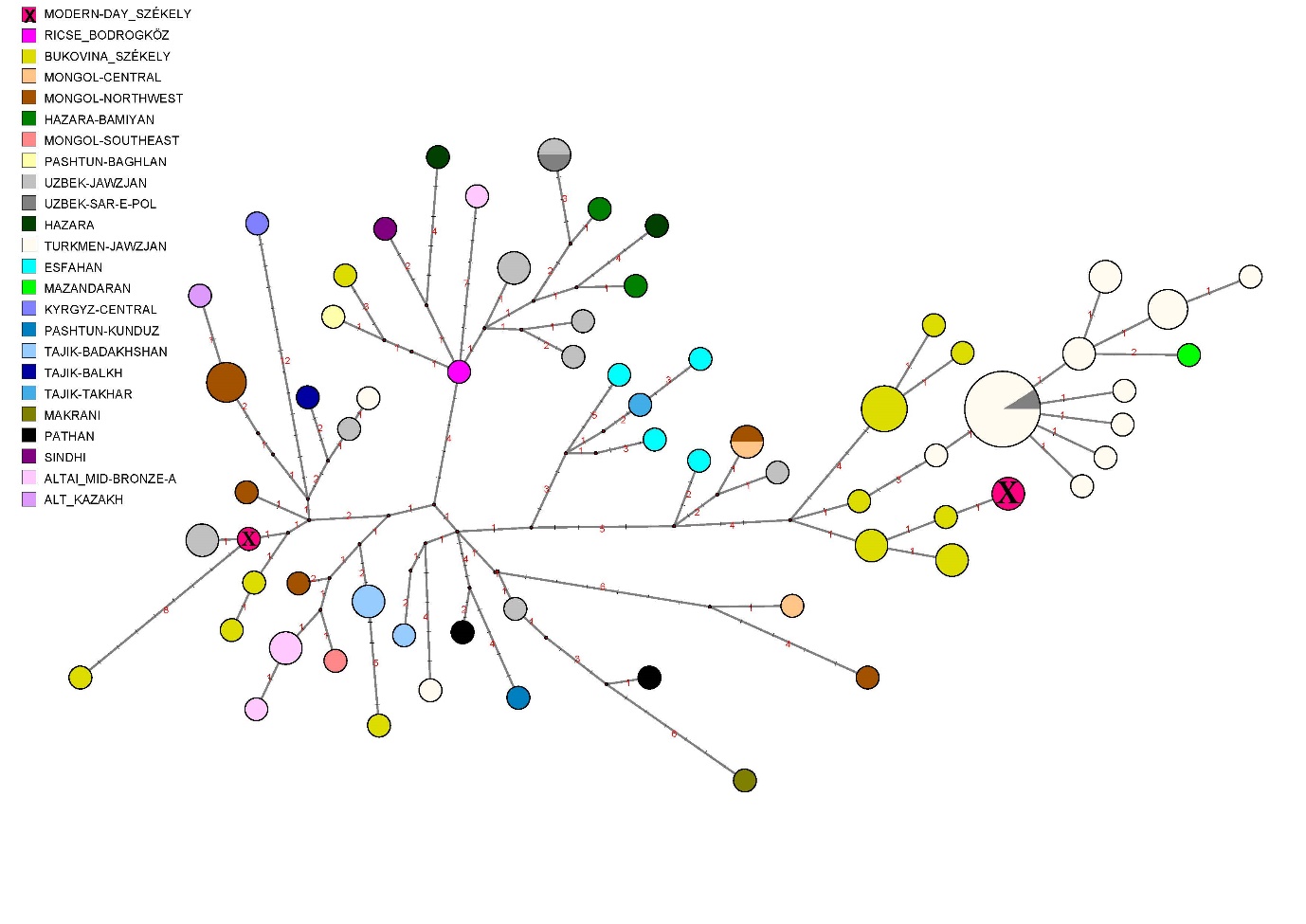
Figure S8: Q- M242 median-joining network based on 16 Y-STRs data. The median-joining network of the haplogroup Q represents populations of diverse geographic origins. The Székely Y chromosome lineages place together with Székely samples of Bukovina.

Figure S9: R1a-M458 median-joining network

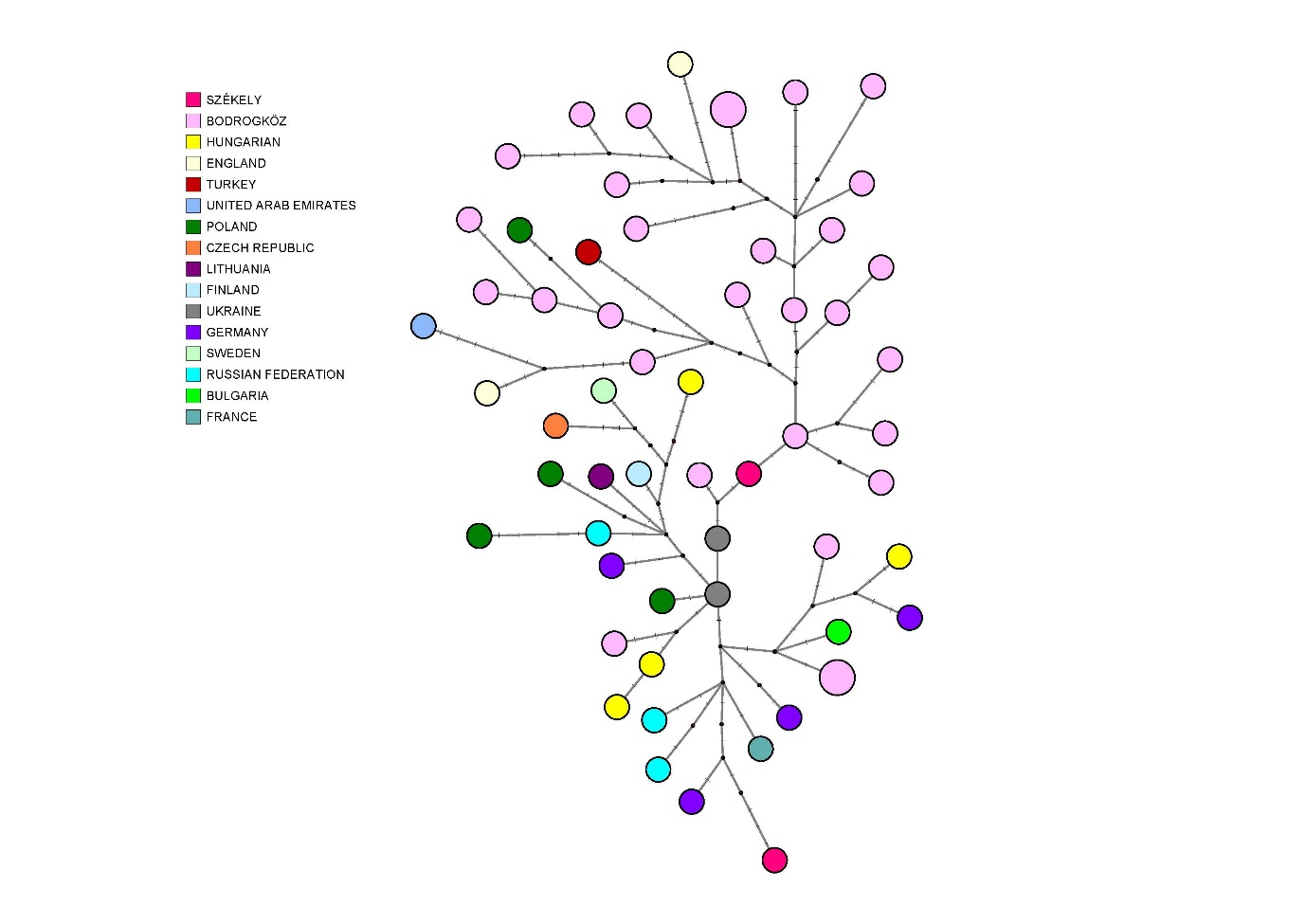

Figure S9: R1a-M458 median-joining network. The analysis was performed based on 23 STRs. We collected data from the public Family Tree Y-DNA database filtering for the subgroup R1a-M458-L1029 and used R1a-M458+ data from Pamjav et al 2017 (Hungarian samples from the Bodrogköz region of northeast Hungary) [6]. This network doesn’t show geographically relevant distribution of the lineages, but shows the divergence of the R1a-M458 types within the Hungarian Bodrogköz population, and the connection of the Székely lineages to some parts of it.

Figure S10: R1a -Z93 median-joining network

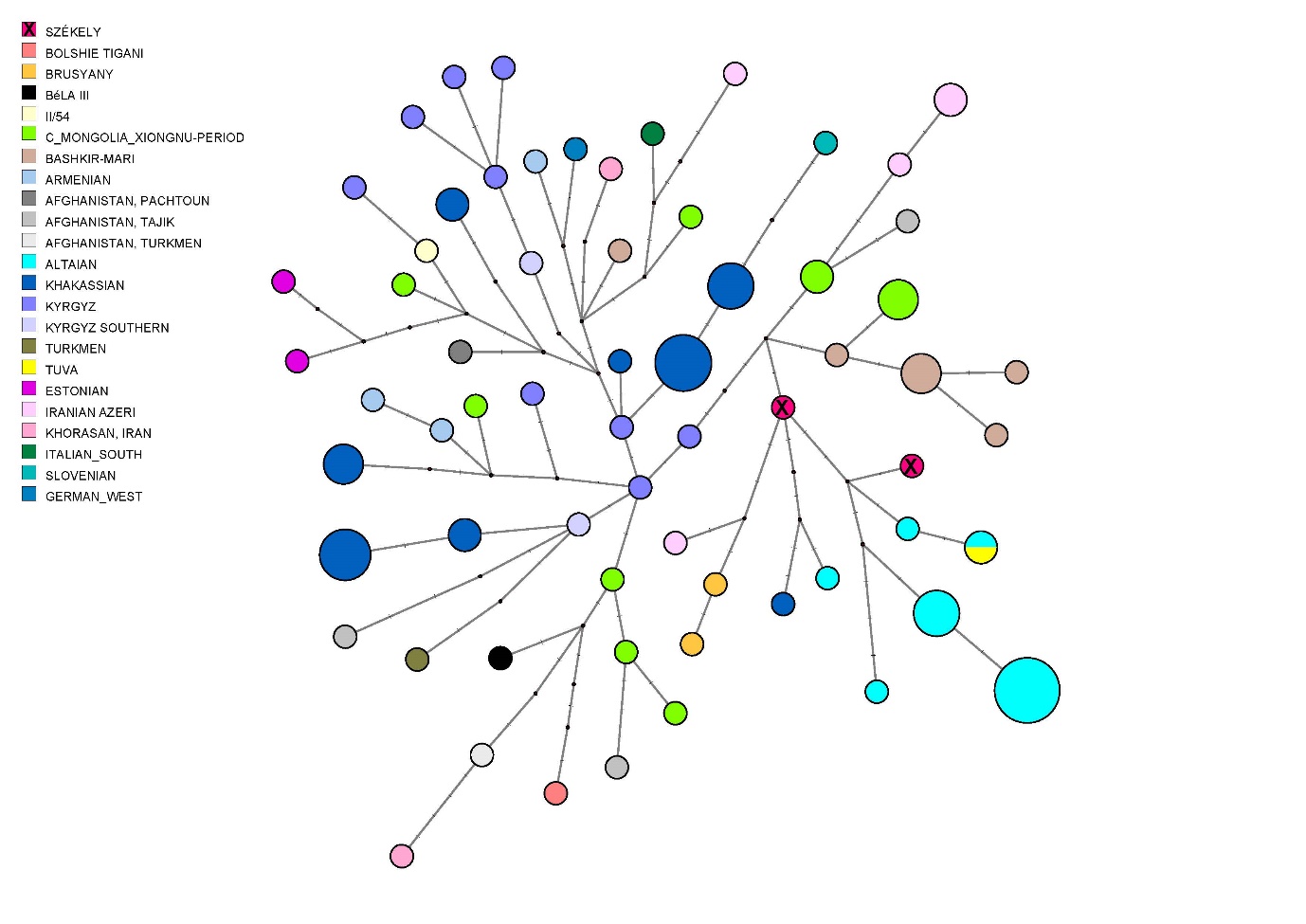

Figure S10: R1a -Z93 median-joining network based on 16 Y-STRs data. For comparison, we collected Y-STR data from the Family Tree Y-DNA database R1a page and filtered the samples for Z93, moreover, we include data from Underhill et al., Olasz et al., Dudás et al., and Szeifert et al. [4,7–9]. Hungarian King Béla III and other skeletal remains originating from the Royal Basilica of Székesfehérvár show a great genetic distance from the Székely samples, just like the Bashkirian Mari males. Samples from Europe (Lithuania, Germany), and Bahrain on the one hand, and Syria, and Kyrgyzstan on the other hand (haplogroup Z2124 - R1a1a1b2a2a (ISOGG v15.73) [10]) are the closest to the two Székely samples respectively.

Figure S11. Multidimensional scaling (MDS) plot

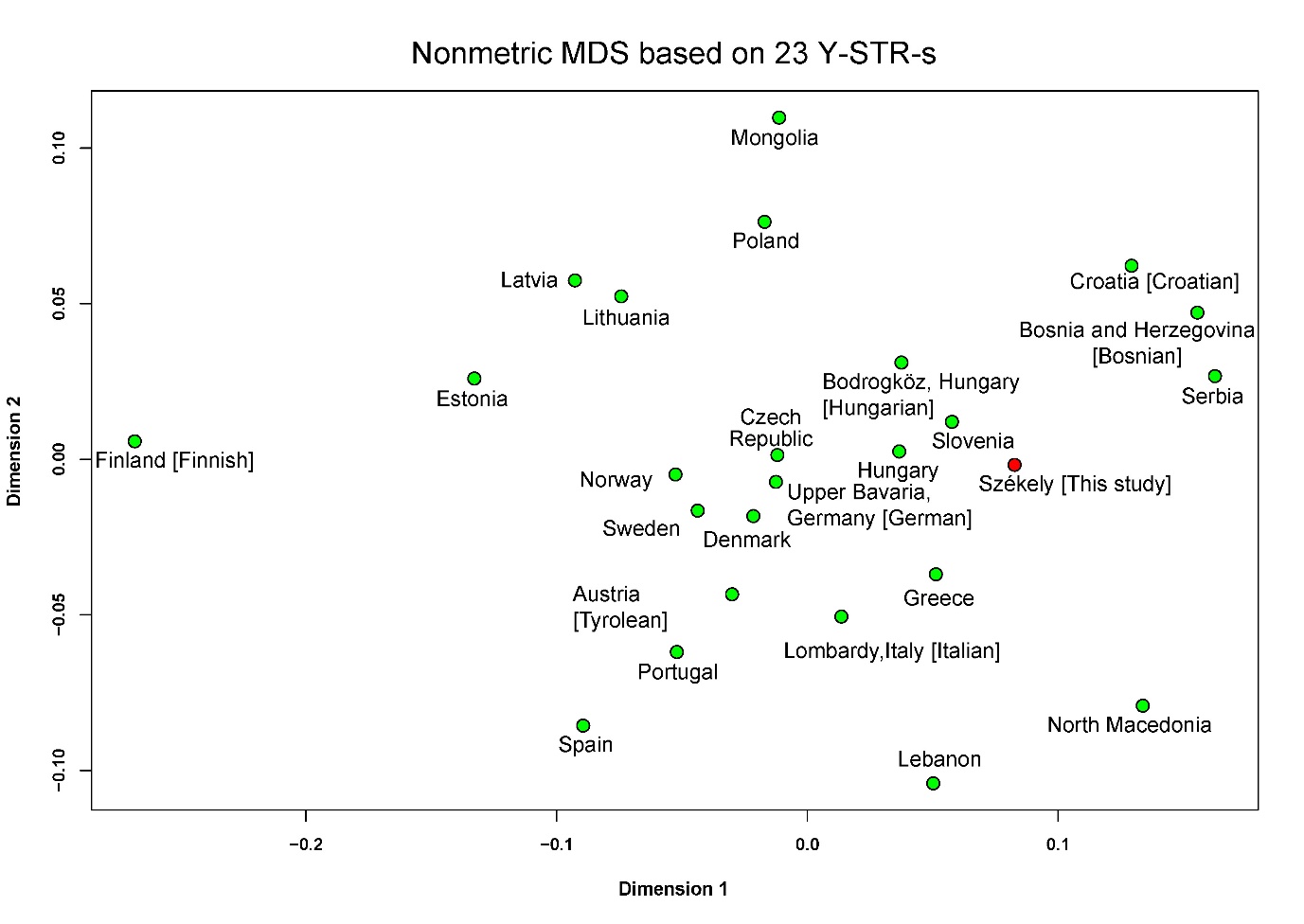

Figure S11. Multidimensional scaling (MDS) plot constructed on R_ST_ genetic distances of 23 STR-based Y haplotype frequencies. The MDS (Kruskals’ non-metric multidimensional scaling) based on 23 Y-STRs (see Supplementary Table S7) shows the strongest paternal genetic connection of the Székelys to the Slovenians (R_ST_ *p*-value is not significant), Hungarians and to Hungarians from the Bodrogköz region (North-East Hungary), and to Greeks. The results of the MDS correspond to the clustering presented in Figure 11. The R_ST_ and *p*-values (see Supplementary Table S6) were generated on the [YHRD website](https://yhrd.org/pages/tools/amova) [11].

*Comparison of the modern-day Székely groups based on HVR-I sequences*

The comparability with reference data from the Carpathian Basin and Romania is rather limited considering that the complete mitochondrial DNA data is missing from the region. We co-analysed the Székelyudvarhely region’s population also sequence-based with previous data on modern-day Székely populations from Csíkszereda [12] and Korond [13]. Given that only Hyper Variable Region I (HVR-I) data are available from these previous, ethnically and geographically relevant studies, we used the HVR-I part of the mitogenomes for the comparison.

The aim of this comparison was to find out whether there is a significant genetic difference between the Székely populations living in different regions of Transylvania.

|  | **Székelyudvarhely region (This study)** | **Csíkszereda (Brandstätter et al. 2007)** [12] | **Korond (Tömöry et al. 2007)** [13] |
| --- | --- | --- | --- |
| n (number of sequences) | 115 | 178 | 76 |
| number of haplotypes (HVR-I region) | 64 | 98 | 50 |
| polimorf sites | 52 | 84 | 54 |
| random match probability | 2.49% | 1.96% | 3.25% |
| mismatch distribution | 4.321 | 4.734 | 4.081 |
| genetic diversity | 0.982 | 0.984 | 0.974 |

**Table 2**. Mitochondrial DNA diversity in three Székely populations based on the sequence data of the HVR-I region (np 16024-16383), excluding the length polymorphisms of the polyC strands.

The genetic diversity is very similar in the three populations, the highest in Csíkszereda. The random match probability (RMP) is a rather high in each case given that we compare relatively small isolated populations [14].

Based on the results of the AMOVA analysis, 98.36% of the total variability between sequence pairs is due to differences within populations and only 1.64% of the total variance can be attributed to differences between populations.

In estimating the significance (p) of F_ST_ values between population pairs, we tested the null hypothesis that population pairs do not differ in their genetic structure. Based on the F_ST_ values, the genetic distance of the three Székely populations can be considered as significant, but the variance between the populations given by the F_ST_ values does not exceed 2% in any of the population pairs (see Table 3).

|  | Korond | Csíkszereda | Székelyudvarhely |
| --- | --- | --- | --- |
| Korond | - | 0.01695* | 0.01523* |
| Csíkszereda | *0.0000* | - | 0.01806* |
| Székelyudvarhely region | *0.0000* | *0.0000* | - |

**Table 3.** Population pairs F_ST_ values (above diagonal) and significance testing (p-value, below the diagonal). * = F_ST_ values observed between population pairs show a significant difference at the p = 0.05 significance level.

Based on mtDNA haplogroup distribution and genetic distance calculation, the previously investigated Székely population from Korond [13] was the most similar to the newly investigated Székely population from the Székelyudvarhely region.

These comparative analyses demonstrate a moderate level of regional variability of the Székelys, the importance of studying multiple Székely regions, and also indicates that by combining the haplogroup data of the three Székely populations we obtain a dataset that can adequately represent the recent Székely population in further comparisons with other populations.
